# Intracellular crowding links dimensionality to cell fate through a mechano-metabolic signalling axis

**DOI:** 10.64898/2026.01.20.700361

**Authors:** M. Meteling, B. van Loo, C. Suurmond, R. Mulders, R. Wijngaarden, K. Govindaraj, L. Geris, I. Meulenbelt, Y.F.M. Ramos, L. Moreira Teixera, J. Leijten

**Affiliations:** Leijten Lab, Department of BioEngineering Technologies, TechMed Centre, University of Twente, Enschede, The Netherlands; Department of Biomedical Data Sciences, Section Molecular Epidemiology, Leiden University Medical Centre, Leiden, The Netherlands; Department of BioEngineering Technologies, TechMed Centre, University of Twente, Enschede, The Netherlands; Prometheus, Division of Skeletal Tissue Engineering, KU Leuven, Leuven, Belgium; Skeletal Biology and Engineering Research Centre, Department of Development and Regeneration, KU Leuven, Leuven, Belgium

## Abstract

Three-dimensional (3D) cell culture systems exhibit more native-like cell behaviour compared to conventional two-dimensional (2D) cell culture systems. However, it remains largely unknown how dimensionality alters cell behaviour. Here, we identify intracellular crowding as a key biophysical parameter altered by cell culture dimensionality, which directs cell fate over a mechano-metabolic axis. Specifically, culture dimensionality controlled intracellular crowding by altering cell volume, which was confirmed across multiple cell types and cell culture platforms. Using chondrogenesis as a model system, we demonstrated that dimensionality-induced intracellular changes lead to improved chondrogenic stem cell differentiation in microtissue culture via a FOXO1 signalling axis. Our findings highlight intracellular crowding as an important parameter of cell culture systems, and present a novel strategy for engineering biomimetic cell culture systems, which has implications for a multitude of applications including disease modelling, *in vitro* drug screening models, developmental biology, and cell-based therapies.

## Introduction

Three-dimensional (3D) cell culture has become essential for a wide range of applications including tissue engineering, disease modelling, drug discovery, and regenerative medicine. For example, 3D cell culture uniquely allows for both studying and modelling inherent 3D processes such as gastrulation [1], organogenesis [2, 3], tumour metastasis [4, 5] and blood vessel formation [6, 7]. In addition, emulating the native cell environment more closely via 3D cell culture, as opposed to two-dimensional (2D) cell culture systems, has been demonstrated to promote more native-like cell behaviour [3, 8–16]. Yet despite its importance and wide-spread use, the fundamental intracellular mechanism(s) that translate dimensionality changes into altered cellular behaviour is currently still poorly understood. Specifically, while it is well established that cell culture platform dimensionality changes alter microenvironmental interactions (e.g. cell-matrix and cell-cell interactions) [8, 9], shift cell metabolism [10–13], and alter mechanotransduction (e.g. YAP/TAZ nuclearization) [14–16], it remains largely unknown how cell culture dimensionality governs these changes in intracellular processes and affects lineage commitment.

## Results

### 3D cell culture induces cell shrinkage and increases intracellular crowding

To investigate how dimensionality drives cell behaviour, we compared cells cultured in monolayers (i.e. 2D) with those self-assembled to microtissues (i.e. 3D) using a microwell array. In contrast to monolayer culture, self-assembled microtissues of expanded primary human adult multipotent stem cells showed rapid and progressive volume shrinkage **(Fig. 1a-b)**, reflected by a marked reduction in cell volume **(Fig. 1c)**. This observation was independently confirmed using flow cytometry **(Fig. 1d).** Further analysis of image-based semi-quantification of cell and microtissue volume showed a clear linear relationship between microtissue and cell volume over time **(Fig. 1e)**. Confocal imaging further revealed that the cellular shrinkage was initiated in the core of the microtissue, and propagated towards its periphery over time, resulting in a near uniform cell volume throughout the microtissue by day six **(Fig. 1f-g).** This is in line with reported computational modelling analysis that predicts increased hydrostatic stress in the core of microtissues as cell adhesion (i.e. cell aggregation) increases, which acts as a compressive force on the cells [17]. Furthermore, we found that the nuclear volume also progressively decreased showing a near-linear relationship with the cell’s volume **(Fig. 1h-i, Fig. S1a)**. Indeed, the inter-nuclei distance notably decreased over time **(Fig. S1b)**. Additionally, cell shrinkage was confirmed in microtissues of various sizes, although the amount of cell shrinkage was aggregate-size dependent **(Fig. 1j, Fig. S1c)**. The slower cell volume decrease in larger microtissues may potentially be explained by shielding effects arising in larger aggregates due to a relatively speaking smaller peripheral region of high lateral tension, which results in reduced compressive forces inside the core, as previously predicted [17]. Thus, this corroborates the previous findings, and implicates increases in intra-tissue hydrostatic pressure as potential driving force for cell shrinkage.

**Fig. 1.**
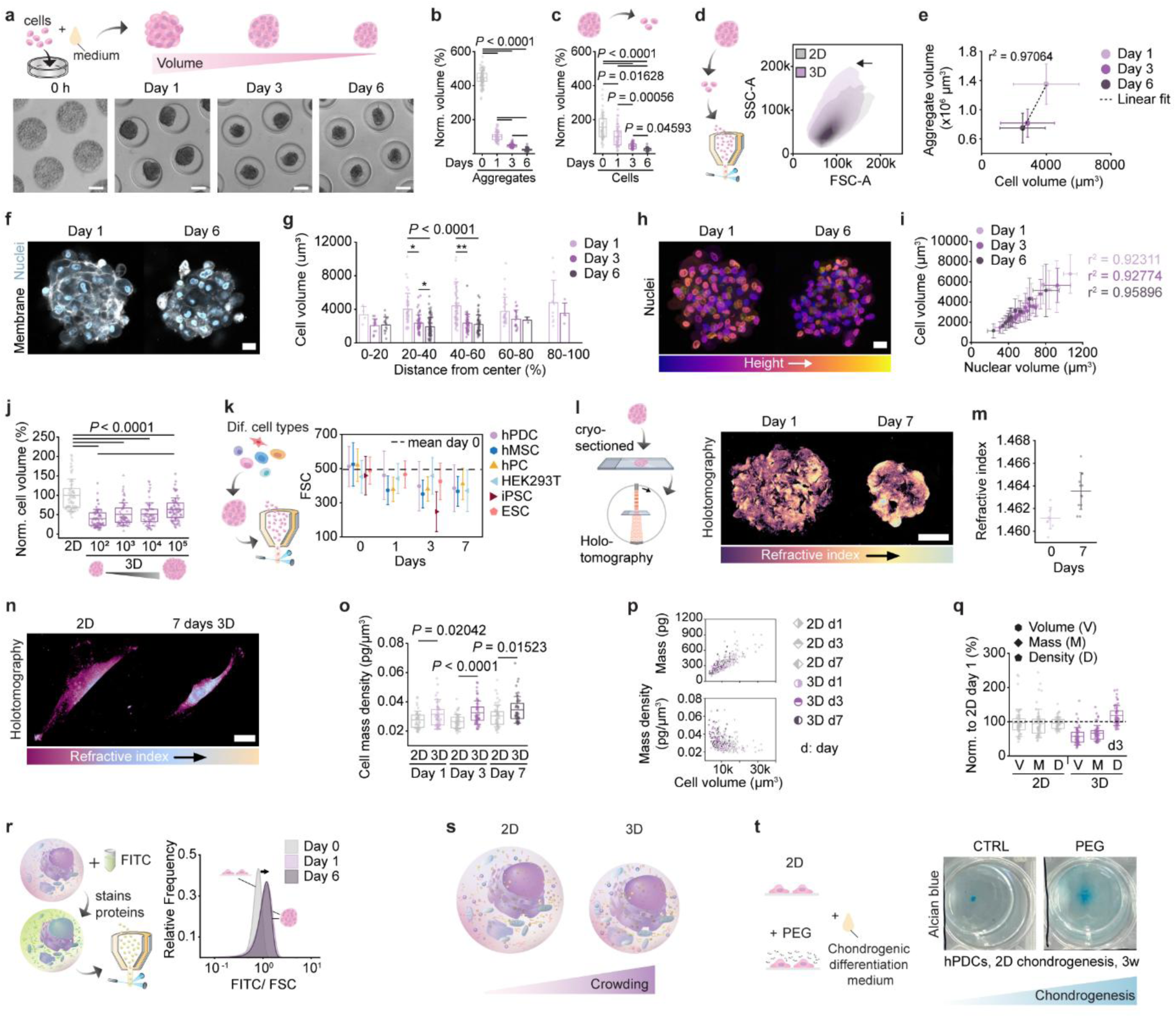
3D cell culture induced cell shrinkage leads to increased intracellular crowding. **a,** Schematic and brightfield images of cell seeding in agarose microwells, and subsequent microtissue formation and shrinkage over time of hPDCs. Scale bar: 100 µm. **b**, Normalized volume of hPDC microtissues and **c**, respective normalized cell volume of microtissue cultured cells over time. Values are based on microtissue or cell diameter measurements from images. **d**, hPDCs volume assessed with flow cytometry following enzymatic harvesting of monolayer cells or dissociation of microtissues following three days in microtissue culture. SSC, side scatter; FSC, forward scatter. **e**, Comparing hPDC microtissue volume to cell volume over time. Values are based on diameter measurements. Linear fitting with respective adjusted r^2^ is shown. **f**, Confocal images of hPDC microtissues after one and six days of culture. Cell membrane (white) and cell nucleus (blue) were fluorescently stained to analyse the *in situ* cell volume. **g,** Respective quantification of the *in situ* cell volume, plotted against the cell’s distance from the microtissue’s centre for one, three and six days in microtissue culture. Scale bar: 20 µm. **h**, Visualization of nuclear compaction in hPDC microtissues over time by spatial colour coding of a fluorescent nuclear staining signal. Scale bar: 20 µm. **i**, *In situ* nuclear volume plotted against cell volume over time of hPDCs cultured in microtissues. **j**, Image based semi-quantification of the cell volume from hPDCs cultured in monolayer or in microtissues of various sizes (10^2^, 10^3^, 10^4^ or 10^5^ cells per microtissue) for three days. **k**, Flow cytometry approximated cell volume (i.e. FSC) of different cell types after zero, one, three or seven days in microtissue culture. **l,** Holotomography images of cryosections from hPDC microtissues that were cultured for one or seven days. Scale bar 20 µm. **m,** Respective quantification of overall refractive index of cryosectioned microtissues. **n**, Holotomography images of hPDCs cultured for seven days in 2D or 3D prior to analysis. Pseudo-colouring indicates differences in refractive index. Scale bar: 20 µm. **o,** Respective quantification of cell mass density of hPDCs cultured for one, three or seven days in either 2D or 3D prior to analysis. **p,** Relation between cell mass and cell volume, and cell mass density and cell volume shown for all conditions. **q,** Normalized cell volume, cell mass and cell mass density of hPDCs cultured for three days (d3) in either 2D or 3D. **r,** Schematic showing FluoCrowd analysis: FITC labelling of intracellular proteins, followed by flow cytometry readout. Estimated crowding (FITC/FSC) of monolayer cultured (day 0) or microtissue cultured (day 1 and 6) hPDCs. **s,** Schematic showing the decreased cell volume and increased in intracellular crowding in 3D cell culture compared to 2D cell culture. **t,** Macroscopic images from representative wells of hPDCs differentiated with or without 1.4% PEG300 supplementation in monolayer for three weeks. Glycosaminoglycan deposition was assessed by Alcian blue staining. Abbreviations: hPDC - human periosteum derived stem cell, hMSC - human mesenchymal stem cell, hPC - human (articular) primary chondrocyte, HEK293T - human embryonic kidney cell line, iPSC - induced pluripotent stem cell, ESC - embryonic stem cell, Norm. – normalized, Dif. – different, FITC **-** Fluorescein Isothiocyanate, PEG – polyethylene glycol. Statistics: The box plots in **b, c, j o, y** give the mean ± 25-75%, with whiskers indicating the S.D.. Statistical significance was assessed using two sample t-tests for comparing 2D to 3D data per time point, using the Welch correction in case of unequal variance (**o**). For multiple comparison test Kruskal-Wallis ANOVA with Dunn post-hoc test was applied (**b, c, g, j**). * p < 0.05, ** p < 0.01.

To exclude the possibility that these observations were cell type-specific, we confirmed that isolated primary cells (human mesenchymal stem cells (hMSCs), human chondrocytes (hPCs)), long-term expanded cell lines (HEK293T (human embryonic kidney 293 cells)), and pluripotent stem cells (embryonic stem cells (ESCs), induced pluripotent stem cells (iPSCs)) also decreased their cellular volume upon 3D microtissue formation **(Fig. 1k)**. Of note, cellular shrinkage proved independent of cell proliferation, which caused microtissue enlargement in fast proliferating cells over time **(Fig. S1d)**. Together, this underscores that self-assembly of cells into 3D microtissues causes condensation and compaction of cells through reduction of cellular volume.

To discriminate whether the observed cell shrinkage was caused by cell-cell contact or by being in a 3D microenvironment, cells were individually encapsulated in a micrometre-thin hydrogel layer, which offers a 3D microenvironment that is void of cell-cell interactions [18, 19]. This confirmed that microencapsulated cells also consistently and progressively decreased in volume over time **(S. Fig. 1e-f)**, although at a slower pace compared to aggregate-cultured cells **(Fig. S1g)**. This indicates that physical constriction of cells imposed by a 3D microenvironment is sufficient to drive cell volume reduction, even when presented by distinct physical surroundings. Taken together, these findings suggest that the elevated dimensionality of 3D cell culture systems, independent of cell type and cell culture platform, elicits a reduction in cell volume.

To investigate what loss of cellular elements associates with the 3D-mediated cell shrinkage, holotomographic imaging was used to simultaneously quantify cell volume and solid dry mass in a spatially resolved manner [20]. This revealed that at microtissue scale, cell volume reduction caused a significant increase in cell mass density **(Fig. 1l-m)**. To evaluate cell mass density at the single cell level, we first tested the reversibility of the 3D-induced cell shrinking by plating one and three-day old microtissues on cell adherent culture plates to induce microtissue melting (i.e. reconfiguration towards a 2D monolayer) **(Fig. S1h)**. Crowding levels in the melted microtissues scaled with microtissue culture duration **(Fig. S1i)**. Given these findings we proceeded to enzymatically disassociate microtissues and reseed the cells on adherent culture plastic. Time-resolved analysis revealed that transitioning back from 3D to 2D associated with a progressive regaining of cell size, although this process operated at a slower pace than when transitioning cells from 2D to 3D culture **(Fig. S1j)**. Holotomographic analysis of microtissue pre-cultured cells confirmed the progressive cell volume decreased over time **(Fig. S1k)**, and indicated elevated intracellular crowding levels as compared to purely monolayer cultured cells, which scaled with 3D cell culture time **(Fig. 1n-o)**. We noted that while the cell mass scaled with cell volume **(Fig. 1p)**, relative changes in cell volume were larger compared to relative changes in cell mass, therefore explaining the increased cell mass density **(Fig. 1q)**. Together, this indicates that 3D cell culture-induced cell volume shrinkage, occurs via water efflux, thus leading to an increase in intracellular crowding.

The increase in intracellular crowding was further verified by using flow cytometry in combination with a total intracellular protein stain for high-throughput single cell analysis. Dividing the total protein stain signal (here FITC fluorescence [21]) by the forward scatter signal, we reasoned that we would obtain a rough estimation of the total protein concentration inside a cell, and thus also a proxy for the intracellular crowding levels. We termed this method FluoCrowd [22]. FluoCrowd analysis confirmed that the aggregate-cultured cells had higher intracellular crowding levels as opposed to the monolayer cultured cells **(Fig. 1r)**, independent of cell type **(Fig. S1l)** or microtissue size **(Fig. S1m)**. Moreover, it was also evident in hydrogel encapsulated cells, which was assessed by confocal fluorescence microscopy **(Fig. S1n)**. Although relatively smaller than the increase observed in aggregates using the same technique **(Fig. S1o)**, it confirmed that the increase in intracellular crowding is not exclusive to scaffold-free 3D cell culture systems. Thus, these data establish increased intracellular crowding as a consequence of dimensionality-driven volumetric shrinkage **(Fig. 1s)**.

To couple the dimensionality-driven changes to cell fate, we used chondrogenesis as a model system as it is a developmental process known to be initiated via tissue condensation [23]. We selected human periosteum derived stem cells (hPDCs) for the cell culture model, as they are the natural cells that drive endochondral bone repair during bone fracture healing [24]. To explore whether increased intracellular crowding could enhance chondrogenesis in 2D, we cultured hPDCs in monolayer in chondrogenic differentiation medium in the presence or absence of an artificial crowder, PEG300. PEG treatment increased glycosaminoglycan deposition visibly at macro level **(Fig. 1u)**. Interestingly, the nuclei size scaled with cell density, being smallest in the areas with the highest cell density in the middle of the well, which corresponded to areas with high glycosaminoglycan deposition **(Fig. S1p-q)**. No statistically significant differences in nuclei area were observed between areas of similar cell densities, comparing artificially more crowded cells to control samples. This links smaller nuclei size, and thus presumably smaller cell volume and increased intracellular crowding, to improved chondrogenesis also in 2D culture. However, we cannot exclude that the increased glycosaminoglycan deposition in the presence of PEG, may also have been facilitated by the more crowded extracellular fluid facilitating matrix deposition [25]. Regardless, taken together, these data hint at a role of intracellular crowding levels in regulating cell fate.

### Crowding-induced metabolic rewiring links dimensionality changes to mitochondrial respiration

Next we investigated whether increased intracellular crowding in microtissues would also associate with altered organellar organization. Transmission electron microscopy revealed major changes in the mitochondrial morphology in microtissues, particularly in the mitochondrial cristae architecture and mitochondrial surface area **(Fig. 2a-c)**. Live-cell confocal microscopy using mitochondrial staining corroborated that cells from microtissues were characterized by higher mitochondrial staining intensities than 2D cultured cells **(Fig. 2d-e),** indicative of mitochondrial crowding. Artificially increasing intracellular crowding in 2D cultured cells using PEG, reproduced the 3D mitochondrial phenotype, indicating that crowding alone is sufficient to drive these changes. Of note, we confirmed that the artificial crowding of monolayer cells was not cytotoxic to the cells, albeit noted that it decreased the cell proliferation rate as is also commonly observed in 3D tissues **(Fig. S2a-c)**.

**Fig. 2.**
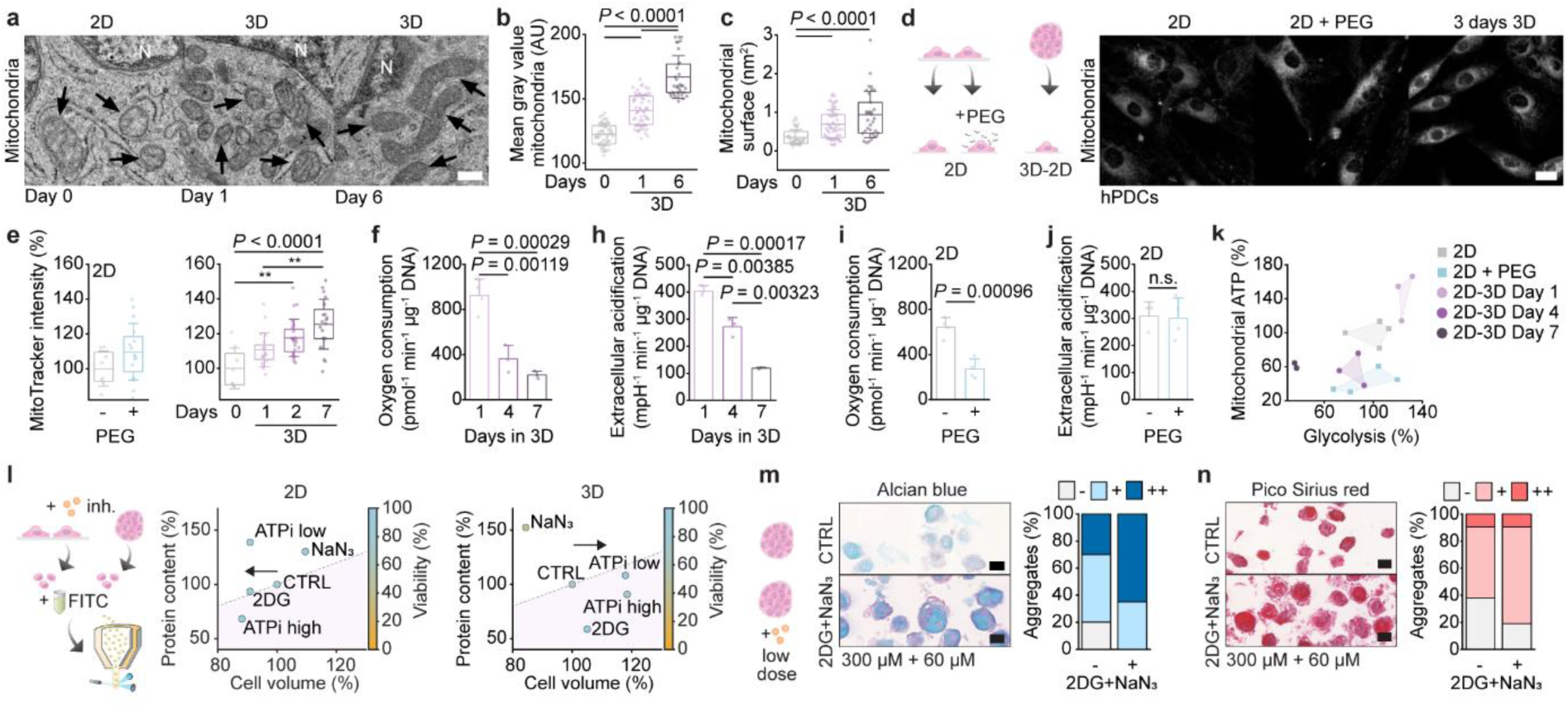
Reciprocal interaction between intracellular crowding levels and mitochondrial respiration. **a,** TEM images of monolayer (2D) and microtissue cultured (3D) hPDCs. Arrows indicate mitochondria. Scale bar: 200 nm. Respective semi-quantification of **b,** the mean gray value and **c,** surface area of the mitochondrial. **d,** Representative confocal fluorescent images of mitochondrial networks in hPDCs cultured in 2D in the presence or absence of PEG, or in microtissues for three days. Scale bar: 20 µm. **e,** Semi-quantification of MitoTracker staining intensity in 2D, or 3D cultured hPDCs over time. **f,** ATP-linked oxygen consumption rate (OCR), and **h,** extracellular acidification rate (ECR) normalized to DNA content per well, of hPDCs cultured in cultured in microtissues for one, four or seven days prior to analysis. **i,** ATP-linked oxygen consumption rate (OCR), and **j,** extracellular acidification rate normalized to DNA content per well, of hPDCs cultured in cultured in monolayer ± 2% v/v PEG300. **k,** Relation of glycolysis-linked ECR to ATP-linked OCR of hPDCs. Data points shown are normalized to the untreated 2D condition. **l,** Flow cytometry approximated crowding of either 2D or 3D cultured hPDCs untreated (CTRL), or treated with NaN_3_ (2 mM), 2DG (50 mM), or a combination of both: ATPi low (2DG 10 mM + NaN_3_ 2mM), ATPi high (2DG 25 mM + NaN_3_ 2mM). Arrow indicates the main trend in cell volume change. Data points represent the normalized median and colour code indicates viability. **m-n,** Three week chondrogenic differentiation of hPDC microtissues in the absence or presence of a low concentration of ATP inhibitors (2DG 300 µM + NaN_3_ 60 µM): **m,** assessed by visualization of glycosaminoglycan matrix deposition (Alcian blue staining) and respective image-based semi-quantification, and **n,** assessed by visualization of collagen matrix deposition (Pico Sirius red staining) and respective image-based semi-quantification (-: < 5%, +: < 50%, ++: >50% of microtissue positively stained). Scale bars: 50 µm. Abbreviations: N – nucleus. 2DG – 2-deoxyglucose, NaN_3_ – sodium azide. PEG – polyethylene glycol. The box plots in **b, c, e** give the mean ± 25-75%, with whiskers indicating the S.D.. Mean ± S.D. is shown in **f, h, i, j**. Statistical significance was assessed using Kruskal-Wallis ANOVA with Dunn post-hoc test (**b, c**) or ANOVA with Bonferroni post-hoc test (**e, f, g, h, i, j**). ** p < 0.01.

As mitochondria are essential for mammalian energy metabolism, and especially for oxidative phosphorylation, we reasoned that 3D-induced cellular shrinkage would also alter a cell’s energy metabolism. Indeed, self-assembly of hPDCs into microtissues significantly decreased mitochondrial respiration and glycolytic rate **(Fig. 2f-h)**. Interestingly, artificially enhanced crowding of monolayer cells emulated the decrease in ATP-linked oxygen consumption rate, but not glycolysis **(Fig. 2i-k)**. However, we noted that the glycolytic reserve of the cells was significantly reduced by PEG addition in 2D, thus also reducing the cells’ glycolytic capacity **(Fig. S2d-g)**. Therefore, increased intracellular crowding seems to inhibit oxidative phosphorylation and to a lesser degree glycolysis, which only becomes visible when the cell needs to compensate for the absence of oxidative phosphorylation. This metabolic dependence on intracellular crowding may be explained by the much higher volume fraction of mitochondria as compared to that of glycolytic enzymes, which can be a limiting factor for mitochondrial ATP production [26]. These results suggest that increased intracellular crowding acts as a biophysical cue rewiring cellular energy metabolism, favouring a lower energy state.

To further explore the relationship between intracellular crowding levels and the cell’s energy metabolism, we inhibited glycolysis and/or oxidative phosphorylation, and subsequently performed FluoCrowd analysis. While changes in cellular volume of monolayer cells in response to glycolytic inhibition or the inhibition of oxidative phosphorylation were in line with previous reports [27], the aggregate-cultured cells surprisingly displayed a near inverse trend **(Fig. 2l)**. While individually 2-deoxyglucose (2DG) (i.e., glycolysis inhibitor) and sodium azide (NaN_3_) (i.e., oxidative phosphorylation inhibitor) had opposite effects on cell volume and intracellular crowding levels, combining these compounds synergistically inhibited 3D-mediated cell shrinkage-induced intracellular crowding in a dose-dependent manner. Together, this suggests a reciprocal interaction between intracellular crowding and mitochondrial respiration, that is governed by cell culture dimensionality. While changes in cellular metabolism in response to microtissue culture have previously been reported [13, 28, 29], we here show that changes in intracellular crowding are a probable cause for a metabolic shift in response to 3D cell culture.

While it is well documented that chondrogenesis changes a cell’s energy metabolism and the cell’s metabolism affects the chondrogenic cell fate [30–32], it less well understood whether 3D-culture induced changes in the cell’s metabolism would alter its chondrogenic capacity. To this end, we chondrogenically differentiated hPDCs under mild inhibition of both glycolysis and oxidative phosphorylation using 2DG and NaN_3_. This demonstrated that inhibition of the energy metabolism, further promoting the 3D culture effect and significantly enhanced chondrogenesis **(Fig. 2m-n, Fig. S2h-o)**. Thus, these findings suggest that introduction of the third dimension via microtissue formation can drive cell fate decisions over a crowding-metabolic signalling axis.

### Actin architecture mediates mechano-metabolic coupling during tissue condensation

To identify the cytoskeletal contributions to dimensionality-driven changes, we exposed hPDC to a variety of cytoskeletal protein inhibitors. This revealed that cytochalasin D (CytD), an inhibitor of actin polymerisation, effectively mitigated microtissue shrinkage **(Fig. 3a-c, Fig. S4a-e)**, while other inhibitors targeting myosin II ATPase or ROCK1/2 mainly affected microtissue shape (i.e. decreased roundness) **(Fig. S3f-g)**, but had minor effects on microtissue shrinkage. Analysing cellular morphology of monolayer cells exposed to the same library of cytoskeletal inhibitors revealed that cell shape was largely maintained in response to each inhibitor, except for CytD **(Fig. S3h),** indicating that the cytoskeleton was largely unchanged. CytD addition caused a rounded up cell morphology, indicating actin depolymerisation. To rule out that CytD treatment affected cell shrinkage due to cytotoxicity, we performed recovery experiments of CytD treated aggregates. Following CytD withdrawal, the aggregates decreased in volume over time, indicating that short-term CytD treatment is reversable and not toxic to the cells **(Fig. S3i)**. Moreover, we noted that letting cells self-assemble to microtissues in the presence of CytD potently inhibited the aggregation process, preventing formation of (spherical) microtissues. Meanwhile, exposing self-assembled microtissues to the same concentration of CytD (only) attenuated microtissue condensation **(Fig. S3j-k)**. Taken together, this indicates that the actin cortex is required for successful formation of microtissues, and is involved in 3D-induced cell shrinkage.

**Fig. 3.**
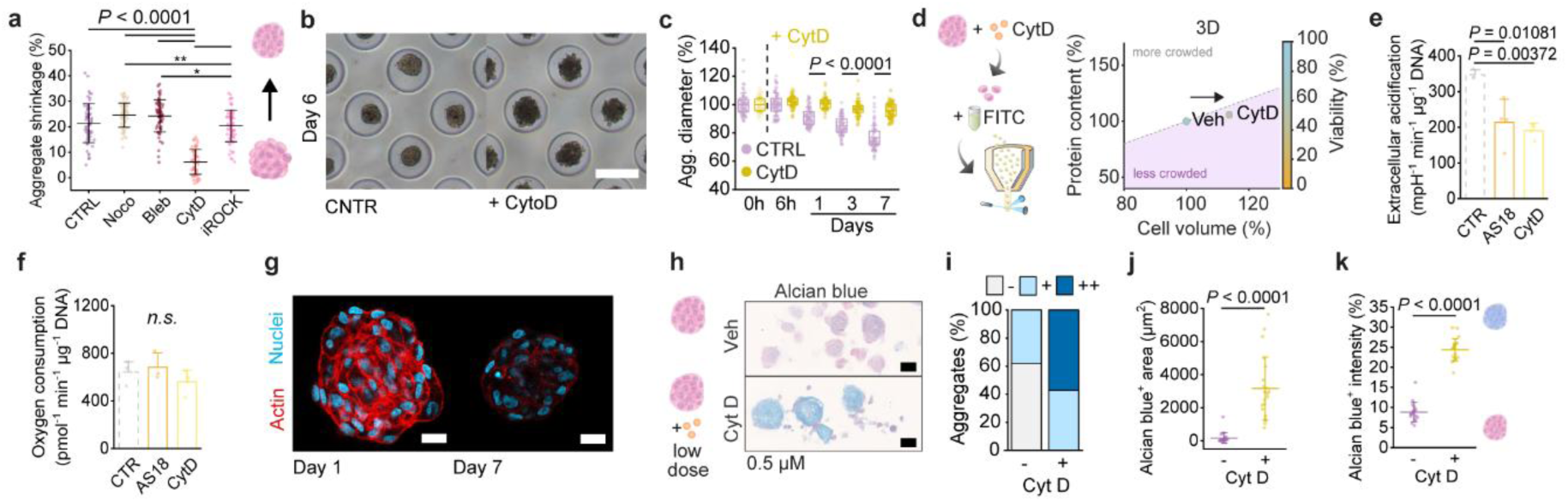
Cytoskeletal remodelling connects physical and metabolic cues in 3D culture. **a,** Image-based semi-quantification of microtissue shrinkage over the course of one week in the presence of different cytoskeletal inhibitors (2.75 µM Noco, 10 µM Bleb, 5 µM CytD, or 10 µM iROCK). **b,** Brightfield images of microtissues cultured with or without CytD (5 µM) at day six. Scale bar: 200 µm. **c,** Image-based semi-quantification of hPDC microtissue diameter over time following CytoD addition (5 µM) six hours after cell seeding. **d,** FluoCrowd analysis of hPDCs with or without CytoD (5 µM) following three days in microtissue culture. Arrow indicates the change in cell volume. Data points represent the normalized median. **e,** Extracellular acidification rate and **f,** ATP-linked oxygen consumption rate of 2D cultured hPDCs in the presence of various inhibitors (1 µM AS18, 1 µm CytD). The control (CTR) values (e, f) are the same as shown in Figure 3j-i respectively. **g,** Representative fluorescence confocal images of actin cortex in hPDC microtissues following one or seven days of culture. A mid-section is shown. Scale bar: 20 µm. **h,** Three-week chondrogenic differentiation of hPDC microtissues with or without low concentrations of CytD (0.5 µM), assessed by visualization of glycosaminoglycan matrix deposition (Alcian blue staining). Scale bar: 50 µm. Respective image-based semi-quantification of: **i,** Alcian blue positive microtissues (-: <5%, +: < 50%, ++: >50% of microtissue positive). n = 21 microtissues per condition, **j,** Alcian blue positive area per microtissue, and **k,** Normalized Alcian blue positive staining intensity per microtissue. Abbreviations: Noco – Nocodazole, Bleb – Blebbistatin, iROCk – Y27632, CytD - cytochalasin D, Veh – vehicle control, AS18 - AS1842856. Statistics: The box plots in **b** give the mean ± 25-75%, with whiskers indicating the S.D.. Mean ± S.D. is shown in **a, e, f, j, k**. Statistical significance was assessed between two conditions using a two sample t-test, and applying Welch correction in case of unequal variance (**c, j, k**). For multiple comparison tests one-way ANOVA with Bonferroni post-hoc test (**e, f**), or Kruskal-Wallis ANOVA with Dunn post-hoc test was used (**a**).

FluoCrowd analysis confirmed that CytD treatment increased the cell volume and slightly decreased intracellular crowding levels in 3D cultured cells **(Fig. 3d)**. A reverse trend for the cell volume was observed for 2D cultured cells, though as the protein content decreased, the intracellular crowding levels decreased as well **(Fig. S3l)**. Thus, this corroborates that the actin cytoskeleton is involved in cell volume regulation, in line with previous reports [33–35], and also affects intracellular crowding levels.

As links between the cellular metabolism and the cells cytoskeleton have recently been reported [36–38], we hypothesized that the dimensionality-driven changes may be mediated via a mechano-metabolic axis. Interestingly, CytD treatment significantly reduced the glycolytic rate but did not affect the ATP linked-mitochondrial respiration **(Fig. 3e-f)**. Notably, in 3D cell culture both 2-DG or CytD addition caused a cell volume increase and crowding decrease, while in 2D cultured cells the respective treatments caused a cell volume decrease. This suggests a reciprocal interaction, which further supports the existence of a mechano-metabolic axis. Following this line of thought, the actin cortex should decrease in microtissue culture over time, given the decrease in glycolytic rate. Indeed, actin fibre staining revealed that the actin cortex decreased over time in microtissue culture **(Fig. 3g)**. Even after enzymatic retrieval and reseeding, this difference persisted, which also confirmed that this observation was not due to dye penetration limitation in microtissues **(Fig. S3m-n)**. This observation may at least in part be explained by previous *in silico* modelling of human progenitor stem cell aggregation [39], reporting that aggregates behave as confluent fluid, which mitigates shear stresses at longer time scales. Moreover, a reduction in cortical stiffness favours compaction [17] and thus presumably condensation, which is in line with our data. In conclusion, these findings position the actin cytoskeleton as a key mediator of the mechano-metabolic axis that drives the dimensionality-induced changes.

To test whether the dimensionality-induced changes in the actin architecture, and by extension the identified mechano-metabolic axis, also affect cell function, we proceeded to differentiate hPDCs in microtissue culture towards the chondrogenic lineage under mild inhibition of the actin cytoskeleton using CytD. Indeed, this enhanced chondrogenesis, as evidenced by a more intense and larger area of Alcian blue positive tissue, and a higher collagen deposition **(Fig. 3h-k, Fig. S3o-p)**. Taken together, this suggests that the dimensionality-induced changes in the actin architecture play a role in regulating cell fate, as demonstrated for chondrogenesis.

### FOXO1 signalling integrates dimensionality-induced changes to regulate cell fate

To gain an understanding of how the mechano-metabolic axis effectuated 3D-induced cell behaviour, RNA sequencing and differential gene expression analysis was performed comparing the gene expression of 2D cultured hPDCs relatively to microtissue cultured hPDCs **(Fig. 4a, Fig. S4a)**. This revealed that 3D culture of hPDCs associated with significantly increased expression of numerous chondrogenesis-associated genes (TGFβ1/2/3, BMP2, SOX4/5, RUNX2), despite the absence of chondrogenic growth factors in the culture medium. Transcription factor target enrichment analysis revealed significant enrichment of SRY-Box Transcription Factor (SOX) 5 and 9 as well as Forkhead box O (FOXO) 1, 3, and 4 **(Fig. 4b)**. While SOX9 as master transcription factor for chondrogenic differentiation confirmed the pro-chondrogenic effects of microtissue culture, the involvement of FOXOs were of particular interest owing to their recently discovered role in inducing chondrogenic commitment of skeletal progenitor cells during endochondral bone healing by acting as an upstream regulator of SOX9 [40]. To determine which FOXO isoform was primarily involved the regulation of the dimensionality-induced effects, we included 3D cultured cells treated with FOXO1 inhibitor (AS1842856) in the analysis. We verified that AS1842856 treatment was not cytotoxic to the cells **(Fig. S4b)**, and successfully inhibited FOXO1 **(Fig. S4c-d)**. This revealed that most of the DE genes were regulated by either FOXO1 or FOXO3, while FOXO4 appeared to exert only a minor role **(Fig. 4c)**. Further focusing on FOXO1/3, a principle component analysis (PCA) revealed two distinct profiles for microtissue and monolayer cultured cells **(Fig. 4d, Fig. S4e)**. Namely, inhibiting FOXO1 in microtissues resulted in a near complete reverse shift towards monolayer levels on the PC2 axis. Further analysis of the top contributors to PC1 and PC2 showed that several of those genes were regulated via FOXO1 or FOXO3 **(Fig. S4f)**. Notably, FOXO1 regulated genes were significantly enriched in both PC1 and PC2, although mainly and most significantly in PC2 **(Fig. 4e)**. Gene ontology-based pathway analysis suggested that PC1 was mostly altered in terms of chemotaxis and extracellular matrix organization, while PC2 genes were mainly involved in developmental processes **(Fig. S4g-h)**, which matches with the known role of FOXO in development [40]. Taken together, these findings suggest that while FOXO1 and FOXO3 are both involved in the gene regulation of DE genes, changes between 2D and 3D gene expression profiles are mainly regulated by transcription factor FOXO1.

**Fig. 4.**
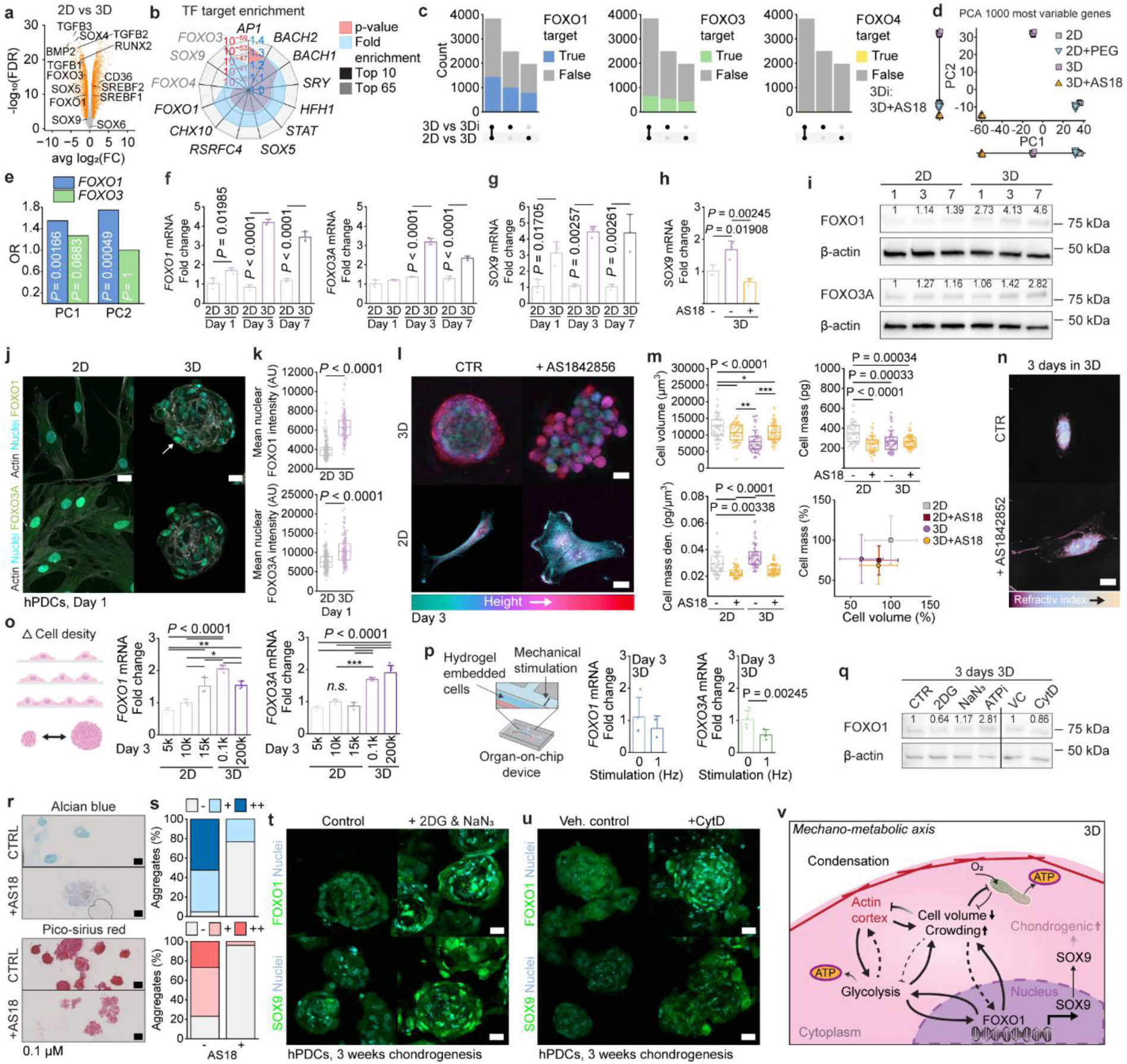
FOXO1 regulates pro-chondrogenic effects of microtissue culture. **a,** Volcano plot showing differentially expressed genes (DEGs; orange) of hPDCs cultured in monolayer or microtissues for three days. **b,** Transcription factor target enrichment of DEGs between monolayer cultured and microtissue cultured hPDCs assessed using DAVID, showing the top 10 enriched genes (based on p-value) and additional FOXOs and SOXs genes (within top 65). **c,** Number of FOXO1/3/4 target genes among the DEGs comparing monolayer to microtissue cultured hPDCs (2D vs 3D), and comparing microtissues to FOXO1 inhibitor (AS18) treated microtissues (3D vs 3Di) or taking the overlapping DEGs of both groups. **d,** PCA plot of the 1000 most variable genes. **e,** Overrepresentation of FOXO1 and FOXO3 regulated genes in PC1 and PC2. **f-g,** Assessment of FOXO1/3 and SOX9 mRNA levels in hPDCs cultured in 2D or 3D for one, three or seven days. Significances were assessed with multiple comparison tests, but are only shown per time point, highlighting the difference between platforms. **h,** SOX9 mRNA levels in responds to FOXO1 inhibition. **i,** FOXO1/3 protein abundance in monolayer and microtissue cultured hPDCs at different time points. **j,** Fluorescence confocal images of hPDCs cultured in monolayer (MIP) or microtissues (mid-section) to assess FOXO1/3 location, and **k**, respective image-based quantification of FOXO1/3 nuclear intensity. White arrow indicates nuclear FOXO1. Scale bar: 20 µm. **l,** Holotomography images of hPDC microtissues and monolayer cells treated with FOXO1 inhibitor (AS18; 1 µM). Maximum intensity projection images are shown with spatial colour coding. Scale bar: 20 µm. **m,** Quantification of cell volume, cell mass and cell mass density of hPDCs cultured in 2D or 3D treated with or without FOXO1 inhibitor (AS18; 1 µM) for three days. Relation between cell mass and cell volume is depicted. **n,** Representative holotomography images (MIPs) with pseudo-colouring for the refractive index of reseeded monolayer and microtissue cultured hPDCs following three days of culture. Scale bar: 20 µm. **o,** FOXO1/3 expression of hPDCs in response to different cell seeding densities in 2D culture, and 3D culture in either a small or large microtissue. **p,** FOXO1/3 expression of hydrogel embedded hPDCs in either static or mechanically actuated culture for three days. **q,** FOXO1 abundance in hPDC microtissues treated with 2DG (50 mM), NaN_3_ (2 mM), ATPi (10 mM 2DG + 2 mM NaN_3_), CytD (1 µM) compared to controls following three days of culture. **n,** FOXO1/3 mRNA levels in responds to different cell densities in 2D and 3D. **r,** Two-week chondrogenesis of hPDC microtissues treated with FOXO1 inhibitor (AS18; 0.1 µM) and untreated control, assessed for glycosaminoglycan content (Alcian blue staining) and collagen deposition (Pico Sirius red staining). Scale bars: 50 µm. **s,** Respective quantification of Alcian blue and Pico Sirius red positive areas in aggregates respectively (-: <5%, +: < 50%, ++: >50% of microtissue positive). n ≥ 15 microtissues per condition. **t,** Three-week chondrogenesis of hPDC microtissues treated with ATP inhibitors (300 µM 2DG + 60 µM NaN_3_) and respective control, assessed for FOXO1 and SOX9 localisation and abundance. Scale bars 20 µm. **u,** Three-week chondrogenesis of hPDC microtissues treated with CytD (0.5 µM) and respective control, assessed for FOXO1 and SOX9 localisation and abundance. Scale bars: 20 µm. **v,** Schematic showing the postulated mechanism for the pro-chondrogenic effects of microtissue culture: mechano-metabolic axis that interfaces with intracellular crowding levels and FOXO1 expression. Dashed arrows indicate suggestive links. Gradient arrow indicates a time-dependence. Abbreviations: AS18 - AS1842856, MIP: maximum intensity projection. Statistics: The box plots in **j, l** give the mean ± 25-75%, with whiskers indicating the S.D.. Mean ± S.D. is shown in **f, h, n, o**. Statistical significance was assessed using a two-sample t-test (**o**), ANOVA with Bonferroni post-hoc test (**f, g, h, n**) or Kruskal-Wallis ANOVA with Dunn post-hoc test (**j, l**). * p < 0.05, ** p < 0.01. Protein abundance was normalized to the β-actin abundance.

Direct comparison of the expressed genes in pristine microtissues and FOXO1 inhibited microtissues also revealed that most chondrogenesis-related genes were significantly less expressed in inhibited microtissues, with the exception of TGFβ3 and RUNX3 **(Fig. S5a)**. It is tempting to speculate that PC1 genes represent the direct changes caused by cellular aggregation, while the PC2 genes are more involved in the downstream effects, leading to a shift in cell fate, mainly governed by FOXO1. Further, for artificially crowded monolayer cells we only noted a small left shift on the PC1 axis towards the microtissues **(Fig. 4d)**, indicating that artificial crowding with PEG (for three days in proliferation medium) recapitulates the microtissue phenotype only marginally. Notably, also FOXO1 levels were significantly less expressed in PEG treated cells as compared to microtissue cultured cells **(Fig. S5a)**. Further analysis using RT-qPCR and western blot, confirmed, that independent of the time point measured, the microtissue-cultured cells, exhibited significantly higher FOXO1 and SOX9 levels, as compared to monolayer cultured cells **(Fig. 4f-g)**. Of note, SOX9 expression was most similar to FOXO1 expression. Pharmacological inhibition of FOXO1 in microtissues also resulted in attenuated levels of SOX9, similar to that in monolayer **(Fig. 4h)**.

Increased FOXO1/3A protein abundance in microtissues further supported the functional relevance of FOXOs **(Fig. 4i)**. We noted that FOXO1 was detectably upregulated on the protein level already after four hours suggesting direct regulation, and its levels progressively continued to increase over the first 24 hours **(Fig. S5b)**. Fluorescence immunostaining also confirmed increased nuclearization of FOXO1/3A in 3D culture **(Fig. 4j-k)**. Notably, unlike FOXO3A, FOXO1 was near exclusively cytoplasmic in 2D cultured cells, thus also supporting the RNAseq findings, identifying FOXO1 as the key regulatory FOXO gene driving cell shrinkage.

While our data suggest that microtissue formation-induced cell shrinkage and increased intracellular crowding result in upregulation of FOXO1, little is known about the involvement of FOXO1 in cell volume regulation. FOXO1 inhibition reduced tissue shrinkage, and led to a granular microtissue morphology by day three **(Fig. 4l)**. Moreover, exposure to the AS1842856 resulted in an increased cell volume and decreased crowding levels in 3D cultured cells, while the opposite trend was observed in 2D cultured hPDCs **(Fig. 4l-n)**. Thus, this indicates that transcriptional activity of FOXO1 plays a role in cell volume and intracellular crowding regulation. This is in line with literature, showing an increased cell volume of chondrocytes in menisci of Col2Cre FoxO1 and FoxO3 one month old knockout mice as compared to wildtype mice [41]. Next, we reduced the cell volume and increased intracellular crowding in monolayer cells by PEG addition, which decreased FOXO1 mRNA and protein levels **(Fig. S5c-f)**. Of note, as compared to the untreated control the PEG treatment decreased cell density, which by itself positively correlated with FOXO1 expression **(Fig. 4o)**, and may thus explain the reduction in FOXO1 levels. Interestingly, FOXO1 expression was slightly elevated in small microtissues (100 cells per aggregate) as compared to larger microtissues (0.2 mil cells). This may link back to differences in cell volume and intracellular crowding levels, but could also suggest cytoskeletal involvement (e.g. differences in cell-cell contact). To investigate if FOXO may be mechano-responsive, we encapsulated individual hPDCs in agarose and subjected them to mechanical stimulation using an organ-on-chip device **(Fig. 4p)**. Both FOXO1 and FOXO3 mRNA expression were attenuated in the mechanically stimulated condition, as compared to unstimulated controls. Notably, it has been reported that FOXO1 expression decreased in human cartilage explants subjected to supraphysiological loading [42]. Addition of CytD to microtissues slightly decreased the protein levels of FOXO1 **(Fig. 4q)**, further supporting the hypothesis that FOXO1 is mechanoresponsive. Therefore, we reasoned that FOXO1 may be involved in regulating the identified mechano-metabolic signalling. Inhibition of FOXO1 resulted in decreased glycolysis **(Fig. 3e)**, while inhibition of glycolysis resulted in lowered FOXO1 protein levels (2DG treatment; **Fig. 4q)**. FOXO1 inhibition did not significantly affect mitochondrial respiration **(Fig. 3f)**, and inhibition of oxidative phosphorylation had only minor effects on FOXO1 protein levels (NaN_3_ treatment; **Fig. 4q)**. Therefore, these findings suggests a reciprocal mechanism between FOXO1 and glycolysis. Together, this links FOXO1 to the identified mechano-metabolic axis. Combined inhibition of both glycolysis and oxidative phosphorylation resulted in a major increase in FOXO1 protein levels (ATPi condition; **Fig. 4q)**. This may explain, why over time FOXO1 expression kept increasing, whereas glycolysis, the actin cortex and mitochondrial respiration decreased. It is however assumed, that this later increase in FOXO1 is caused via a different pathway, and less dependent on the cytoskeletal activity.

To corroborate whether FOXO1 also facilitates chondrogenesis, we chondrogenically differentiated hPDCs in the presence of low concentrations of the pharmacological FOXO1 inhibitor. Indeed, FOXO1 inhibition, blocked chondrogenic differentiation of hPDCs microtissues **(Fig. 4r-s)**. This is an interesting finding, as FOXO1 has been found to be relevant in bone healing in mice by regulating the formation of cartilage tissue [40]. Thus, FOXO1 plays a pro-chondrogenic role [43]. Notably, FOXO1 and SOX9 nuclear protein levels were elevated in hPDCs aggregates cultured under mild ATP inhibition **(Fig. 4t)**, and this was also observed for the aggregates treated with low concentrations of CytD **(Fig. 4u)**. Taken together, these findings establish FOXO1 as a potential key transcription factor linking dimensionality-induced changes in intracellular processes to chondrogenesis **(Fig. 4v)**.

## Discussion

Our findings infer that dimensionality directs cell fate in a mechano-metabolic manner, which is guided by alternations in cell volume and intracellular crowding. While dimensionality has long been recognized as a determinant for cell behaviour, the intracellular mechanism orchestrating the translation of the physical context into transcriptional programs remained elusive. Using chondrogenesis as a model system, we demonstrated that following the transition from 2D to 3D cell culture, cells shrink in volume via water efflux, inducing changes in intracellular crowding that orchestrate cytoskeletal remodelling, metabolic rewiring, and FOXO1-driven transcriptional programs, which enhance lineage commitment. The identified mechano-metabolic axis in conjunction with intracellular crowding, provides a conceptual framework for understanding the enhanced physiological relevance of 3D cell culture systems. This expands our current understanding of how changes in cell energy metabolism and actin cytoskeleton jointly influence chondrogenesis [30, 44, 45].

The decrease in cell volume in 3D cell culture was found to be neither cell type nor 3D cell culture platform specific, suggesting that our findings may have broader implications. During embryonic development chondrogenesis is initiated via mesenchymal condensation, an event characterized by cell clustering and increased cell density [46]. As such, microtissue culture, unlike monolayer cell culture, may allow for this process to occur *in vitro*. Thus, this study may potentially also provide new insights into cell condensation processes occurring during embryonic development, identifying intracellular crowding as a potential key biophysical parameter orchestrating cell fate. Notably, increased cell density has been reported to cause volumetric compression in intestinal organoids, locally increasing intracellular crowding levels and regulating intestinal stem cell fate [47]. In addition, a recent paper on embryonic trunk development revealed that the end-stage cell phenotype is controlled early on during development by the balance between glycolysis and oxidative phosphorylation [48]. Future studies should explore if the mechano-metabolic regulation in conjunction with intracellular crowding plays a role in directing cell fate during organogenesis and tissue formation preceded by cell condensation.

Understanding how intracellular crowding orchestrates cell fate opens new opportunities for optimising existing 3D cell culture systems, including disease models and engineered living constructs. By altering intracellular crowding levels, the cell’s perception of its microenvironment is likely changed, which has also been reported for artificially crowded 2D cultured cells [22]. Notably, our findings hint at the existence of an ideal window for optimal chondrogenic differentiation, which likely involves a lower energy metabolism and a reduced actin cortex in 3D culture. Future work should investigate whether intracellular crowding can be tuned as a design parameter to control cell fate decisions, and thus (more) predictably engineer living constructs, which would be beneficial for tissue engineering and regenerative medicine applications alike.

## Methods

### Cell culture

Human periosteum derived stem cells (hPDCs) were isolated from the periosteum of three donors as previously described [49]. The cells were obtained with ethical approval by the local ethical board and written consent by the patients. The isolated cells were pooled, and initial cell expansion (till passage 6) was performed at KU Leuven. For further expansion, cells were cultured in proliferation medium consisting of DMEM high glucose (4.5 g L d-glucose) supplemented with 10% v/v fetal bovine serum (FBS) (Sigma) and 1% v/v Penicillin/Streptomycin (Pen/Strep) (ThermoFisher). Cells were subcultured prior to reaching about 80% confluency using 1x TrypLE (Gibco^TM^, ThermoFisher). Medium was refreshed three times a week. For the chondrogenic differentiation experiments, cells were used up to passage 12.

Human chondrocytes (hPCs) were isolated from human cartilage samples. The cartilage samples were obtained from osteoarthritis patients who underwent total knee replacement surgery. The use of patient material was approved by the local ethical committee and written consent was obtained from all patients. Cartilage pieces were extracted from macroscopically non-degenerated areas of the joint and incubated with collagenase II in DMEM under agitation at 37 °C overnight. Following recovery of hPCs, the cells were directly frozen. For cell expansion, cells were cultured in chondrocyte medium consisting of DMEM high glucose supplemented with 10% v/v FBS, 1% v/v Pen/Strep, 0.2 mM ascorbic acid 2-phosphate (ASAP; Sigma Aldrich), 0.35 mM L-Proline (Sigma Aldrich), and 1x non-essential amino acids (NEAA; ThermoFisher). When reaching about 80% confluency, cells were passaged using 0.25% Trypsin-EDTA. Medium was refreshed three times a week.

Human bone marrow mesenchymal stem cells (hMSCs) were isolated from fresh human bone marrow samples as previously described [50]. Cells were obtained with ethical consent given by the local ethical committee and written consent of the patient (METC\06025). The cells were expanded in Minimum Essential Medium-α containing nucleosides (α-MEM, Gibco™, ThermoFisher) supplemented with 1% v/v ASAP and 1% v/v GlutaMAX™ (100 X, Gibco™, ThermoFisher)). Prior to reaching confluency the cells were subcultured using 0.25% Trypsin-EDTA. Medium was refreshed three times a week.

HEK293T is a human embryonic kidney cell line (HEK293). Cells were cultured in DMEM high glucose medium with 1% Pen/Strep and 10% FBS. Passaging was done using 0.25% Trypsin-EDTA. Medium was refreshed three times a week. For the one week microtissue culture the medium was refreshed up to two times daily.

Induced pluripotent stem cells (iPSCs) and human embryonic stem cells (hESCs) were cultured in vitronectin (Gibco™, ThermoFisher) coated 6-well plates in well-defined Essential 8™ Medium (Basal medium plus Supplement^TM^; Gibco™, ThermoFisher). For dissociation, the cells were shortly incubate with 1 mL EDTA (0.5 mM in DPBS, 37°C, 4 min), followed by aspiration. Cells were removed from the plate by carefully resuspending them in 1 ml of E8 medium. The medium was refreshed on a daily basis. When seeding the cells in the microwells Y-27623 (10 µM) was added to the cell culture medium of the iPSCs.

All cells were maintained in culture at 37 °C in a humidified air atmosphere at 5% CO_2_.

### Microwell fabrication for 3D cell culture

Agarose microwells were fabricated with the use of a PDMS molds as previously described [51]. Briefly, for creating PDMS molds, first a patterned silicon wafer was fabricated using photolithography. An elastomeric stamp was subsequently obtained by PDMS replica molding. The resulting stamp design consisted of round pillars of either 200 µm diameter with a interpillar distance of 100 µm for aggregates of about 100 cells, or 400 µm pillar diameter with 200 µm inter-pillar distance for aggregates of 1000 cells. For aggregates of 10,000 cells, PDMS molds were created by casting PDMS on a plastic base plate with pillars, of 2.2 mm height, 1 mm diameter, and an inter-pillar distance of 1.5 mm. All stamps were designed to fit in a 12 well plate. For creating agarose microwell inserts, PDMS molds were placed with the pattern facing upwards in 6-well culture plates, and fully covered with 3% w/v agarose solution dissolved in sterile PBS (Ca^2+^/Mg^2+^ free). Following gelation at 4 °C, the PDMS molds were carefully removed, and agarose microwell inserts were created by punching out disks containing the microwell pattern with raised edges severing as walls. Following transfer of the agarose microwell inserts to a 12 well culture plate, sterile PBS was added, and the plate was centrifuged at 500 g for 5 min. The agarose microwell inserts were fabricated under sterile conditions. Additionally, the inserts were sterilized by 15 min of ultra-violet light exposure prior to use. Following UV treatment and a few hours prior to cell seeding, the microwell inserts were washed with PBS and transferred to cell culture medium. The plates were again centrifuged, and medium was refreshed, to remove any remaining PBS. The plates were placed in the incubator at 37°C and 5% CO_2_ till the cells were seeded.

### 3D cell culture: microtissue formation

For microtissue formation, agarose microwell inserts were first covered each with a small volume of cell culture medium (around 0.5 ml) containing the desired cell concentration. Cell concentration was adjusted to total microwell number per insert and required cell number per microtissue. The cells were given about 10 min to sediment into the microwells, prior to centrifugation of the well plate at 500 g for 3 min. Following centrifugation, additional medium was carefully added, to a final volume of 1.5 ml per well. The medium was refreshed thrice a week.

For pellet culture (0.1-0.2 mil cells), the cells were seeded in 15 ml conical tubes and centrifuged at 500 g for 5 min. The cap was not completely screwed tight to allow for gas exchange. The medium was exchange three times a week as well.

### 3D cell culture: cell encapsulation

Dextran-tyramine (Dex-TA) was synthesized as previously described [52–54]. The microfluidic chips for droplet formation and crosslinking were prepared using standard soft lithography techniques, as previously reported [18]. For single cell encapsulation, hPDCs were grown till about 80% confluency and subsequently harvested. The resulting cell suspension was strained using a 40 µm cell strainer to obtain a pure single cell suspension. Single cell encapsulation was performed as previously described [18]. In short, a cell-laden hydrogel precursor solution was prepared containing a cell concentration of 3.33 mil cells, 10% Dex-TA, 8.33% OptiPrep (Sigma Aldrich) and 42.5 U ml^-1^ horseradish peroxidase (HRP; Sigma Aldrich) in PBS. A flow focusing droplet generator with a channel height of 28 μm was used for droplet generation. Flow rates of 9 μl min^-1^ for the oil phase, and 1.5 μl min^-1^ for the cell-laden hydrogel precursor solution were used. A H_2_O_2_ diffusion-based crosslinking chip connected to the droplet generator, was employed for delayed enzymatic crosslinking of the Dex-TA. Encapsulated cells were recovered by washing with Pico-Break solution in the presence of PBS. The obtained single-cell-laden microgels were cultured for up to one week in hPDC proliferation medium.

### Inhibition assays

Inhibitors were either dissolved in DMSO (cytochalasin D (CytD) (Enzo Life Sciences), blebbistatin (Enzo Life Sciences), nocodazole (Enzo Life Sciences), Y-27623 (Enzo Life Sciences), AS1842856 (Calbiochem), EIPA (5-(N-Ethyl-N-isopropyl)amiloride; Enzo Life Sciences), NBBP (5-nitro-2-(3-phenylpropylamino)-benzoic acid; Enzo Life Sciences), oligomycin A (Cayman Chemical), FCCP (Carbonyl cyanide 4-(trifluoromethoxy)phenylhydrazone; Sigma Aldrich), Rotenone (Sigma Aldrich), Antimycin A (Sigma Aldrich)) or in cell culture water (gladolium III chloride (Enzo Life Sciences), ouabain (Enzo Life Sciences), 2-deoxyglucose (Sigma Aldrich), sodium azide (extra pure, Technipur®; Merck)). The stock solutions were aliquoted and stored in the -30°C or -80°C freezer. The inhibitors were added to either complete proliferation medium, or chondrogenic differentiation medium at the concentrations specified in the figure legends and/or text. For DMSO-dissolved inhibitors the added inhibitor solution volume to the cell culture medium was not above 0.1% (v/v) and for water-dissolved inhibitors below 1% (v/v). Unless otherwise specified, the inhibitors were added following cell attachment in 2D culture, or cellular self-assembly to microtissues (3D culture). Artificially enhanced crowding of monolayer cells was achieved via the addition of 2% (v/v) Poly(ethylene glycol) (PEG) 300 (BioUltra, Sigma Aldrich) directly to the cell culture medium.

### Image-based microtissue and cell size analysis

To determine microtissue size, images were taken from the aggregated cells inside the microwells using a brightfield microscope (EVOS or Nikon) with a 4x objective. ImageJ was used to determine microtissue (ferret) diameter and area. To this end either thresholding was applied, or alternatively images were opened in Adobe Photoshop, an additional transparent image layer was created, and aggregates outlines were manually traced. The image of the outlines was saved and imported to ImageJ for analysis. The microtissue volume was calculated based on the microtissue diameter, assuming that the aggregates have a spherical shape. A similar procedure was employed to do a semi-quantification of the cell size. Dissociated aggregates and trypsinized monolayer cells, were imaged in suspension directly after detachment, while still exhibiting a round cell shape. The same microscopes were used with a 10x objective. The cell (feret) diameter was determine using ImageJ, and used to calculate the cellular volume, again assuming a spherical shape.

### Viability staining

Cell viability was assessed by staining with 4 µM Calcium AM (Sigma Aldrich/ Merck) and 2 µM Ethidium homodimer (Sigma Aldrich/ Merck). Aggregates were incubated for 20-30 min at room temperature, while monolayer cells were incubated for 10 min. Encapsulated cells were incubated with 2 µM Calcein AM and 4 µM Ethidium homodimer in PBS for 15 min. Imaging was performed with an EVOS microscope using GFP and RFP filter cubes for the Calcein AM and Ethidium homodimer signal, respectively. ImageJ was used for image analysis. The cell counter plug-in was employed to determine the number of live and dead cells.

### Nuclear, cell membrane and actin staining

Fixated cells (10% formalin for 15 min at RT), were stained with DAPI (ThermoFisher) and Alexa-488-phalloidin staining (ThermoFisher) to visualise the cell nucleus and actin fibres respectively. After washing with PBS twice, cells were permeabilised in 0.1 % v/v Triton-X100 (in PBS; Sigma) solution for 4 min, and again washed twice with PBS. The microaggregates or monolayer cells were incubated in phalloidin solution according to manufacturer’s instruction at RT for 45 min. Two washing steps with PBS were followed by counterstaining with DAPI (1:100 in 1% (w/v) bovine serum albumin (BSA; Sigma)). Monolayer cells were incubated for 5 min, and aggregates for 10-15 min in DAPI staining solution. Subsequently, the samples were washed thrice with PBS and imaged with a confocal microscope (Zeiss LSM880) or a home-built two-photon laser scanning microscope. Both are part of the BioImaging Centre at the University of Twente.

Live cell staining was performed using DRAQ5 (ThermoFisher) and Cell MaskTM green (ThermoFisher) for nuclear and cell membrane staining respectively. After washing twice with PBS, aggregates were incubated in cell mask green solution (dilution to 1:1000 in PBS) for 15 min at 37°C. Subsequently, the aggregates were washed twice with PBS and stained with DRAQ5 (5 µM) for 30 min at 37°C. The aggregates were imaged directly afterwards at 37°C and 5% CO_2_ (microscope stage incubator) using a confocal microscope (Nikon confocal A1). For acquisition z-stacks were taken with a 20x objective.

### Flow cytometry and FluoCrowd analysis

Monolayer cells were dissociated from the well plates or cell culture flasks using either 0.25% Trypsin-EDTA, 0.5 mM EDTA (in DPBS) or 1x TrypLE treatment, depending on the cell type. Microaggregates were harvested from the microwells by thorough resuspension in medium. The aggregates were dispersed via incubation in either 0.25% Trypsin-EDTA, 0.5 mM EDTA (in DPBS) or 1x TrypLE solution. The obtained cell suspensions were strained with a cell strainer of 40 μm. Following washing in PBS, cells were either directly stained or first fixated in excess of 10% formalin solution for 10 min at room temperature. Total intracellular protein staining was performed by incubating the cells in 5 µg/ml FITC staining solution (freshly diluted from stock solution (20 mg/ml) in PBS) at a cell concentration of no more than 1.2 mil cell ml^-1^. Fixated cells were incubated at 4°C for 30 min, while living cells were incubated at 37°C for 20 min. After incubation the cells were washed with PBS twice and then resuspended in a final volume of 100-200 μl PBS for flow cytometry analysis (around 1 mil cells ml^-1^ cell density). The stained cells were kept on ice or in the fridge until analysis. For dead cell detection, additional samples were prepared. Designated samples were resuspended in Propidium iodide (Sigma-Aldrich, P4864) staining solution (10 µg ml^-1^ in PBS) in a 1:1 ratio with sample volume and incubated for 2 min on ice immediately before flow cytometry analysis. Unstained control samples were taken along as well.

Flow cytometry analysis was performed either on a FACSCaliburTM (BD Biosciences) or a FACS Aria 2 (BD Biosciences). For each condition a minimum of 20000 events was obtained. The instruments were operated with BD CellQuestTM (BD Biosciences) or BD FACSDivaTM (BD Biosciences) software respectively. Collected data was further analysed using either Flowing Software 2 (version 2.5.1) [55] or Matlab (MathWorks). For the latter, the Flow cytometer GUI [56] in combination with fca_readfcs [57], both available online at MATLAB central file exchange, were utilized.

### Mitochondrial staining

For mitochondrial image cells were harvested from monolayer or microtissue conditions the day prior to analysis and seeded in ibidi dished for high resolution imaging. Overnight incubation allowed the cells to attach. The next day the cells were incubated with MitoTracker^TM^ green (ThermoFisher) solution (1 µM) for 10 min, followed by an additional 25 min in combination with DRAQ5 (5 µM) solution. The cells were washed once with pre-warmed complete medium. For artificially increasing the crowding in the monolayer cells, 2% (v/v) PEG300 was added to the cell culture medium about 30 min prior to imaging. Live-cell imaging was done using a confocal microscope (Zeiss LSM880) with a 40x water-immersion objective (1.2NA) at 37°C and 5% CO_2_ using a stage incubation chamber. Z-stacks were taken from all conditions with z-steps of 1 µm. Semi-quantification of MitoTracker staining intensity was performed using ImageJ.

### TEM

Samples were fixated using 2.5% glutaraldehyde and 2% paraformaldehyde in 0.1M Na-cacodylate buffer calibrated at pH 7.4 at room temperature for 2 hours, after which the samples were washed in 0.1M Na-cacodylate buffer. Post-fixation was then performed using an aqueous solution containing 1% osmium tetroxide for 2 hours, after which the samples were dehydrated using a graded ethanol series, and stained using uranyl acetate dissolved in 70% ethanol. Samples were subsequently infiltrated using a 1:1 mixture composed of epoxy resin (Agar100) and propylene oxide at 4°C for 60 minutes, followed by an overnight incubation using a 2:1 mixture of agar and propylene oxide, after which the samples were embedded and polymerized in 100% agar for two days. Subsequently, sample were cut in 50–60 nm sections using a Leica UCT ultramicrotome. Cut samples were stained with uranyl acetate for 10 minutes, and lead citrate for 3 minutes. The resulting stained sections of the samples were photographed using a transmission electron microscope JEOL JEM-1400 equipped with a 2k x 2k Veleta (Olympus SIS, Tokyo, Japan) camera. Mitochondrial morphology was semi-quantified via artisan measurements in ImageJ.

### Metabolic assays

The day prior to metabolic analysis, hPDC aggregates and monolayer cells were harvested and dispersed to a single cell suspension using 1x TrypLE. The cells were subsequently seeded in a Seahorse XF96 microplate (Agilent, 101085004) at a seeding density of 6000 cells per well. Following cell attachment, the next day the proliferation medium was replaced with assay medium consisting of RPMI (Sigma, R1383) supplemented with 2 mM GlutaMax (Gibco, 35050-061) and 5% FBS (Sigma, S0615). For the Mito stress test, 10 mM glucose (Gibco, 15023021) was additionally added. Medium osmolarity was adjusted to either 300 mOsm or 450 mOsm using PEG300. Mito stress test and glycolytic test were performed on a metabolic flux analyser (Seahorse Bioscience, North Billerica), according to manufacturer’s instructions. For each condition at least three replicates were taken along. Prior to each assay the plates were incubated at 37°C in a non-CO_2_ incubator for one hour. In short, for the Mito stress test, oligomycin (1 µM), FCCP (3 μM), and rotenone (1 µM) combined with antimycin (1 µM) were injected. The Glycolytic stress test involved injections of glucose (10 mM), oligomycin (1 μM) and 2-DG (10 mM). Both extracellular acidification rate (ECAR) and the oxygen consumption rate (OCR) were recorded. Following each injection, three measurements were taken. The resulting data was normalized to cell numbers based on DNA quantity per well as determine afterwards (QuantiFluor dsDNA system, Promega, E2670).

### Holotomography

Live-cell imaging at 37°C with 5% CO_2_ was performed for monolayer or microtissue cultured hPDCs. Following one, three or seven days of culture, the cells were harvested from cell culture flasks or microwells and seeded in ibidi high-resolution imaging dishes. Following cell attachment overnight, the cells were imaged in the morning. Aggregates seeded in ibidi dishes were given approximately 20h for attachment and melting prior to imaging. Cells were kept inside a cell culture incubator at 37°C and 5% CO_2_ till imaging. Cell volume and cell mass analysis was done using the TomoCube LipidAnalyser software. Thresholding was chosen such that the entire cell was detected. Cell mass density was calculated by dividing the cell mass by the cells volume. Pseudo-colouring of the refractive index or height data, was performed in Fiji (ImageJ).

For holotomographic imaging of tissue sections, cryosections of aggregates were used. In short, sectioned tissue was transferred to super frost glass slides, hydrated in PBS, air dried and mounted using aqueous mounting medium. Imaging was performed at room temperature. Pseudo-colouring of the refractive index was performed in Fiji (ImageJ).

### Mechanical stimulation

The organ-on-chip devices for mechanical stimulation were prepared as previously described [58]. The chips were seeded with hPDCs in a 2% w/v agarose (UltraPure Low Melting Point agarose, Invitrogen) solution (dissolved in PBS) at a concentration of 1.5 mil cells ml^-1^. One minute after injection hPDC proliferation medium was added to the medium channel. Chip designs with a single mechanical actuation chamber orientated in parallel to the hydrogel chamber were used. Mechanical actuation of the bulk hydrogel was performed at 300 mbar for 1h as previously described [58]. One day after cell seeding in the chips, the stimulation was started, and continued for three subsequent days in a row. As controls, cells were seeded in the same device but received no mechanical stimulation. On the last day of stimulation the hydrogels were extracted from the chips, the cells were lysed and RNA extraction and RT-qPCR analysis were performed.

### Total RNA extraction and RT-qPCR analysis

Monolayer, microtissue or pellet cultured hPDCs in proliferation medium were lysed and RNA was isolated using the NucleoSpin RNA isolation kit (Macherey-Nagel) as per manufacturers instruction. For agarose embedded hPDCs, cell lysis and RNA isolation were performed using the RNeasy Micro kit (Qiagen). cDNA synthesis was performed using the iScript cDNA synthesis kit (BioRad). For Reverse transcription with quantitative PCR (RT–qPCR) the SensiMix kit from Bioline (GC, Biotech) was used with SYBR Green dye. RT-qPCR was run on a Bio-Rad CFX96 device (Bio-Rad). The 2^-ΔΔCt^ method was used to analyse the gene expression. Normalisation of the gene expression was done with the expression of the housekeeping gene *RPL13A*. *FOXO1* (NM_002015): 5’-AGGGTTAGTGAGCAGGTTACAC-3’ (forward), 5’-TGCTGCCAAGTCTGACGAAA-3’ (reverse) published by Song et al. [59], *FOXO3* (accession nr. NM_001455): 5′-AGGACAGAACCGTGCATAGG-3′ (forward), 5′-ATCACCCTGACTCAGAACCG-3′ (reverse) published by Osswald et al. [60], *SOX9*: 5’-TGGGCAAGCTCTGGAGACTTC-3’ (forward), 5’-ATCCGGGTGGTCCTTCTTGTG-3’ (reverse) from Harvard primer data bank, *RPL13A* (accession nr. NM_012423): 5’-AAAAAGCGGATGGTGGTTC-3’ (forward), 5’-CTTCCGGTAGTGGATCTTGG- 3’ (reverse) published by Pombo-Suarez et al. [61].

### RNAseq

hPDCs were cultured in monolayer (6-well plates) or microtissues (microwells) in proliferation medium for three days. Microtissues of about 100 cells each were prepared. For artificial crowding, 2% v/v PEG300 was added to monolayer cell culture medium. FOXO1 inhibition was done by AS1842856 (1 µM) supplementation. For each of the four conditions, seven replicates were prepared. For RNA extraction, the monolayer cells were directly lysed in the well plate. Microtissues were first harvest from the microwell inserted, washed and then lysed. RNA isolation was performed using the NucleoSpin RNA isolation kit (Macherey-Nagel). The RNA purity and concentration were tested using a Nanodrop 2000. The samples were transported on dry ice to Macrogen Europe (The Netherlands) for RNAseq analysis. Prior to RNAseq analysis the samples underwent quality control again. The RNA sequencing library was prepared using a TruSeq stranded RNA library prep kit (Illumina) and sequencing was performed on Illumina NovaSeq 6000 platform (40M reads). Reads were assessed with FastQC (v0.12.0) [62], aligned to reference genome by STAR (v2.7.10b) [63], adapter trimmed by cutadapt (v4.4) in Python (v3.11) [64], assessed with Samtools (v1.17) [65, 66] and Picard (v3.0.0) [67], and converted to count matrix by HTSeq(v2.02) in Python [68]. Differential Expression Analysis (DEA) was performed in R (v3.6.3) using edgeR (v3.28.1) [69] and counts were normalized to counts per million (CPM), selection criteria of a false discovery rate below 0.001 and a fold change higher than 1.5 were chosen. Transcription factor target enrichment analysis was performed on the differentially expressed genes between monolayer and microtissue cultured hPDCs using the Functional Annotation Tool of the Database for Annotation, Visualization, and Integrated Discovery (DAVID) [70, 71]. Further analysis was performed in R (v4.1.0). Experimentally validated(chromatin immunoprecipitation) FOXO targets in the Gene Transcription Regulation Database (GTRD) (v21.12) [72] was used for determining FOXO target genes within the differentially expressed gene sets. To determine the 1000 most variable gene expression by VST-expression, the RNA expression counts were first normalized using variance stabilising normalisation with Deseq2 (v1.32.0) [73], and then coefficient of variance ranking was performed using multiClust (v1.22.0) [74]. Principle component analysis (PCA) was carried out on the 1000 most variable features of each condition. The genes contributing for more than 0.05% to PC1 (586 genes) or PC2 (420 genes) were considered significant, and were assessed for DAVID-defined FOXO1/3 target enrichment by Fisher Exact test. Gene ontology overrepresentation analysis was performed on the significant contributors to PC1 and PC2 for Gene Ontology (GO) terms biological processes (BP) using clusterProfiler (v4.0.5) [75]. Sematic similarity correction was performed to simplify outcomes based on 70% sematic overlap.

### Western blot

Following cell culture, cells or aggregates were washed twice with PBS. Lysis was performed on ice using RIPA lysis buffer. Following successful lysis, the lysate was centrifuged at 14000 g at 4 ⁰C for 15 min. The supernatant was transferred to vials, snap-frozen on dry ice and stored at -80 ⁰C till further analysis. A BCA assay (Thermo Scientific) according to manufacturer’s instructions was performed to determine protein concentration. SDS-PAGE gels (Bio-Rad) were loaded with 10 µg of protein per condition, and a protein standard (BioRad, Precision Plus Protein^TM^) was taken along. Following protein separation, the blot was transferred to a membrane using the Trans-Blot transfer pack (BioRad) and the TransBlot Turbo device (BioRad). Membranes were washes in TBS-T (0.1% Tween-20 (Sigma)), then blocked in 5% BSA in TBS-T for 1 hour at RT under dynamic conditions. For FOXO1/3A staining, the membranes were incubated with rabbit anti-FOXO1 (1:1000; Cell Signaling Technology, no. 2880), or rabbit anti-FOXO3A (1:1000; Cell Signaling Technology, no. 2497) in 5% BSA in TBS-T under dynamic conditions, at 4 ⁰C overnight. As loading control, the membrane were stained with mouse anti-beta-actin (1:2000, Abcam, ab8224) in 5% BSA in TBS-T under dynamic conditions, at 4 ⁰C overnight. Subsequently, membranes were washed in TBS-T buffer under dynamic conditions, prior to incubation with HRP-conjugated secondary antibody (DAKO). To this end, membranes were incubated with the respective secondary antibody (1:2000) in TBS-T buffer for 1 h under dynamic conditions. Repeating the washing under dynamic conditions, the targeted protein signal was detected by chemiluminescence using the SuperSignal or FemtoSignal HRP substrate kit following manufacturer’s instructions. Imaging was done on a FluorChem M imager (ProteinSimple). Band detection and signal intensity were obtained from the imager software or determined using ImageJ.

### Immunocytochemistry

For immunostaining, hPDCs were seeded in ibidi dishes for high-resolution imaging following culture in either monolayer (cell culture flasks) or aggregates for up to one week. Incubation overnight allowed for cell attachment. The next day the cells were fixed with 4% paraformaldehyde. For immunostaining of entire aggregates, the aggregates were collected from the microwells, washed with PBS and fixed with 4% paraformaldehyde as well. Following fixation, the samples were permeabilized with 0.5% Triton-X100 in PBS, and blocked in 5% BSA plus 5% goat serum in PBS for one hour. Aggregates were incubated during all incubation steps under dynamic conditions. Overnight incubation at 4 ⁰C with the primary antibody FOXO1 (1:100; Cell Signaling Technology, no. 2880) or FOXO3a (1:100; Cell Signaling Technology, no. 2497) in blocking solution with 0.5% Tween-20 followed. The next day the sample were washed thrice with 0.5% Tween-20 PBS solution. Secondary antibody staining was done with anti-rabbit goat Alexa Fluor 488 (1:500) in 5% BSA with 0.5% Tween-20 for 2h at room temperature. Next, the samples were washed again three times with 0.5% Tween-20 PBS solution. Nuclear counterstaining was performed with DAPI (1:100 in 1% BSA) for 5 min for monolayer samples and 10 min for microtissue samples. The samples were washed in PBS. As control 2^nd^ AB only stained samples were taken along. Imaging was performed with a confocal microscope (Zeiss LSM880 or Nikon A1) using either a 20x or 40x (air/water-immersion) objective. Z-stacks were taken from both monolayer and microtissue cells.

### Differentiation assay

For chondrogenic differentiation hPDCs were seeded in microwells (about 100 cells) and transferred to chondrogenic differentiation medium consisting of DMEM high glucose supplemented with 1x Insulin-Transferrin-Selenium (ITS), 0.2 mM ASAP, 0.35mM L-Prol, 1x Sodium pyruvate and 1% v/v Pen/Strep. In addition 10 ng ml^-1^ of recombinant human TGFβ-3 and 10^-7^ M dexamethasone were freshly added on the day of medium refreshment. For the emulation studies the respective inhibitors were freshly added to the medium in end concentrations as reported in the figures. Thrice a week the medium was refreshed. Cells were culture for up to three weeks.

For chondrogenic differentiation in 2D in the presence of an osmotic agent, hPDCs were seeded in 12-well plates at a high seeding density (50k per well). The next day, the wells were transferred to chondrogenic differentiation medium with or without 1.4% v/v PEG300. Medium was refreshed three times a week. The cells were kept in culture for 21 days.

### (Immuno)histochemistry

Following culture in chondrogenic differentiation medium, the samples were collected, washed and fixed using 4% paraformaldehyde. Following permeabilization with 0.1% Tween-20 in PBS, samples were incubated with 15% sucrose solution. After sedimentation the samples were transferred to 30% sucrose solution at 4 ⁰C incubated overnight. Samples were embedded in cryomatrix (Epredia) and snape-frozen in isopentane (stored at -80 ⁰C). Cryosections of 12 µm were cut following standard procedure and using a cryotome. Cut sections were directly transferred to superfrost glass slides (ThermoFisher) and left to dry at RT. Samples were either used for immunostaining or histology staining.

For immunostaining the sections were hydrated in PBS, then permeabilized with PBS-Tween-20. Sections were blocked with 10% goat serum plus 2% BSA for 1h. Primary antibody incubation was done in 5% BSA with 0.5% Tween-20 using FOXO1 (1:100), FOXO3A (1:100) or SOX9 (1:100) for 1 h. Next, sections were washed thrice with 0.5% Tween-20 in PBS and twice with 5% BSA. Subsequently, sections were incubated with the anti-rabbit goat Alexa Fluor 488 (1:500; ThermoFisher, cat. 10 236 882). The same washing steps were repeated. Nuclear counter staining was done with DAPI solution (1:500 in PBS) for 5 min. Samples were washed three times with PBS, then transferred to tap water. Sections were shortly dried, then mounted with fluorescence mounting medium (DAKO). Imaging was performed with a confocal microscope (Zeiss LSM880) using a 20x objective. Z-stacks were taken from all conditions.

To determine the glycosaminoglycan content sections were stained with Alcian blue (Sigma). For visualisation of collagen deposition, sections were stained with Pico Sirius red. Briefly, following hydration in demiwater, sections were incubated in Alcian Blue (pH 1) for 30 min. Sections were washed with tap water and counterstained with Nuclear fast red for 5 min. Sections were washed again, shortly dried and mounted with aqueous mounting medium (Agilent Dako). For Pico Sirius red staining, following hydration, the sections were first counterstained in Weigert’s Hematoxylin solution for 8 min. Sections were washed with DI water, and incubated for 2 min in Phosphomolybdic acid (Polysciences). Following another rinse in DI water, sections were incubated for 1 h in Picosirius red stain (Polysciences), then directly transferred to 0.1 N hydrochloride acid solution (Polysciences) and incubated for 2 min. Following dehydration, the samples were cleared, and mounted with GLC mounting medium (Sakura). Imaging of both stainings was performed using a NanoZoomer (Hamamatsu).

### Visualisation & Statistical analysis

Data analysis and plotting was done in Matlab, R-studio, or Origin Pro. Except for the RNAseq data, the statistics analysis was performed using Origin Pro. In most cases the individual data points are included in the graphs. The mean and standard deviation are provided unless otherwise specified in the figure legend. Statistical significance was assessed by two-tailed student t-tests (with Welch correction if necessary), one-way ANOVA with Bonferroni post hoc test, or Kruskal-Wallis ANOVA with Dunn post hoc test (as specified in figure legends). A difference was considered statistically significant for *P* < 0.05. Schematics were created using Adobe Illustrator, except for the schematic of a confocal microscope and a microscope glass slide which were both obtained from BioRender.

## Acknowledgements

We would like to thank A. van der Ham for expert support with the metabolic assays.

