## Supporting information for "Intracellular crowding links dimensionality to cell fate through a mechano-metabolic signalling axis"

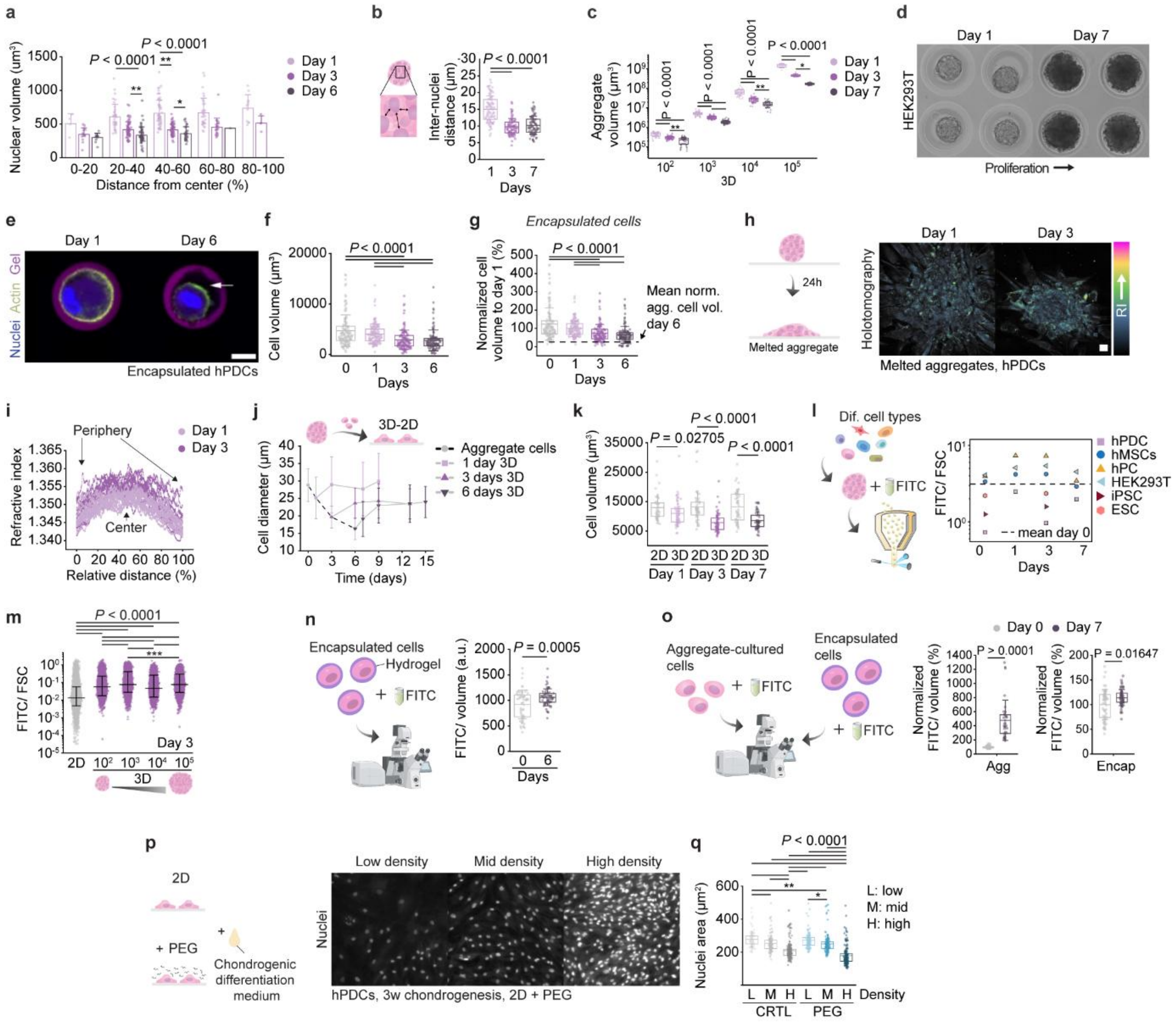


**Fig. S1 | a,** In situ nuclear volume of cells inside an microtissue based on confocal z-stack images. Nuclear volume is plotted against the cell's distance from the microtissue's centre for one, three, and six days in microtissue culture. **b**, Inter-nuclei distances inside microtissues assessed based on two-photon fluorescence imaging of microtissues. **c,** Microtissue volume shrinkage over time displayed for microtissues of different sizes. **d,** Cellular microtissues of HEK293T cells after one and seven days of culture, revealing cell proliferation. **e,** Encapsulated hPDCs in Dex-TA microgels, and **f,** respective quantification of the cell volume over time. Scale bar: 10 µm. **g**, Normalized cell volume of microtissue cultured and hydrogel encapsulated hPDCs on the day of encapsulation (0 days) and after one, three, and six days of culture. **h,** Holotomography images of melted microtissues following one or three days of microtissue culture. Pseudo-colouring based on refractive index (RI). **i,** Semi-quantification of refractive index across melted microtissues. Scale bar: 20 µm. **j,** Cell diameter of hPDCs cultured in microtissues for one, three, or six days, and subsequently reseeded in monolayer (3D-2D cells) and tracked over time. Dots indicate mean values ± S.D.. **k,** Holotomography based calculated cell volume of hPDCs cultured in either 2D and 3D for one, three, or seven days prior to analysis. **l,** FluoCrowd analysis of various cell types after zero, one, three or seven days in microtissue culture. Dots represent mean values. **m,** FluoCrowd analysis of hPDCs cultured in monolayer (2D) or microtissues of various sizes ($10^2$, $10^3$, $10^4$ or $10^5$ cells per microtissue) for three days. Black lines indicate median ± 5-95% range. **n,** Confocal-based estimation of crowding levels using the FITC staining, of Dex-TA hydrogel encapsulated hPDCs. **o**, Normalized FITC fluorescence signal divided by cell volume (i.e. FSC) for hPDCs cultured either in microtissues or encapsulated in Dex-TA hydrogel for zero to six days. For day zero, encapsulated cells are analysed after encapsulation, while for the microtissue-cultured cells, suspension cells (2D) were analysed. **p,** Visualisation of nuclei in chondrogenically differentiated hPDCs in 2D in the presence of PEG300 for three weeks. Shown are areas of low, medium, and high cell density for artificially crowded cells, and **q,**

respective semi-quantification of nuclei area in control and PEG treated wells. The box plots in **b, c, f, g, k, n, o, q** give the mean ± 25-75% , with whiskers indicating the S.D.. The bar graphs indicate the mean ± S.D.. Statistical significance was assessed using two sample t-tests for comparing 2D to 3D data per time point, using the Welch correction in case of unequal variance (**n**), or using a two-tailed Mann-Whitney test (**o**). For multiple comparison test one-way ANOVA or Kruskal-Wallis ANOVA with Bonferroni post-hoc test or Dunn post-hoc test respectively was applied (**b, c, f, g, k, m, q**). * $p < 0.05$, ** $p < 0.01$.

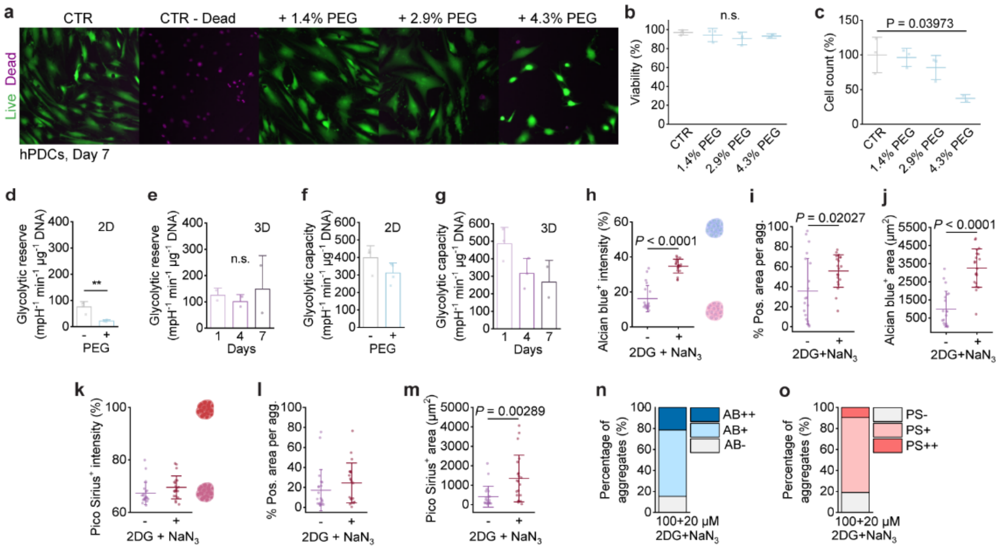


**Fig. S2 | a,** Representative fluorescent images of hPDCs cultured in the presence or absence of various PEG concentrations (1.4-4.3% v/v PEG 300) for seven days, and subsequently stained with Calcein AM (live, green) and Ethidium homodimer (dead, pink). A positive control for cell death (CTR - Dead) was included. **b,** Respective quantification of cell viability (percentage live cells) and **c,** cell count following seven days of culture are shown. Initial seeding density was the same for all conditions. **d-e,** Glycolytic reserve of hPDCs cultured either in **d,** 2D with or without 2% v/v PEG300, or **e,** 3D prior to analysis for one, four, or seven days. **f-g,** Glycolytic capacity of hPDCs cultured either in **d,** 2D with or without 2% v/v PEG300, or **e,** 3D prior to analysis for one, four, or seven days. Readout from glycolysis stress test (**d-g**). **h-j,** Semi-quantification of glycosaminoglycan deposition (Alcian blue staining) of hPDC microtissues cultured in chondrogenic differentiation medium under mild ATP inhibition (300 µm 2DG and 60 µm $NaN_3$) for three weeks. **k-m,** Semi-quantification of collagen deposition (Pico Sirius red staining) of hPDC microtissues cultured in chondrogenic differentiation medium under mild ATP inhibition (300 µm 2DG and 60 µm $NaN_3$) for three weeks. **n,** Three week chondrogenic differentiation of hPDC under weak ATP inhibition (100 µm 2DG and 20 µm $NaN_3$) assessed by Alcian blue (glycoaminoglycan deposition) and **o,** Pico Sirius red (collagen deposition) staining. Semi-quantification of the percentage of microtissues negative (<5% positive), mildly positive (< 50% positive), and very positive (>50% positive) for the respective staining is depicted. Abbreviations: 2DG – 2-deoxyglucose, $NaN_3$ – sodium azide. AB – Alcian blue, PS – Pico Sirius red. Statistics: Mean ± S.D. is shown in **b-m**. Statistical significance was assessed between two conditions using a two sample t-test, for unequal variance the Welch correction was applied (**d, f, h-m**). For multiple comparison tests ANOVA with Bonferroni post-hoc test (**e, g**), or Kruskal-Wallis ANOVA with Dunn post-hoc test was used (**b, c**).** $p < 0.01$.

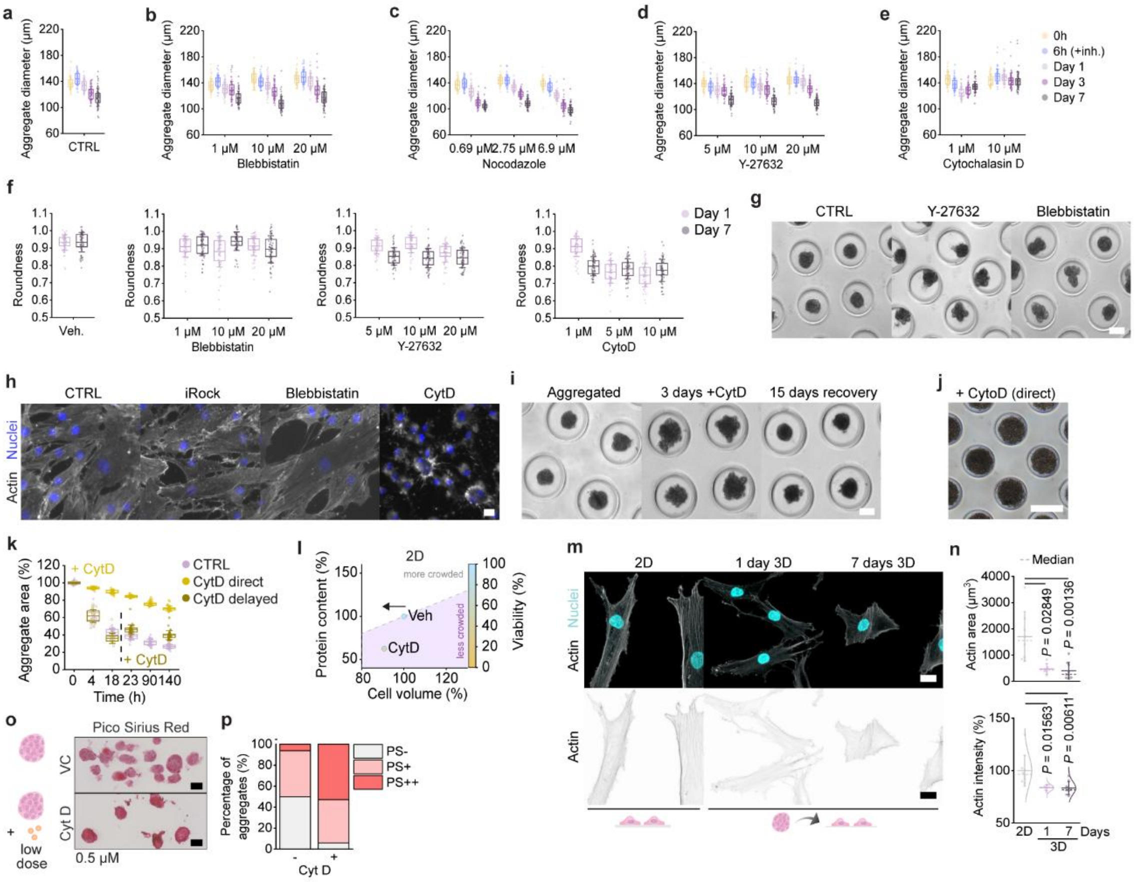


**Fig. S3 | a-e,** Time-resolved semi-quantitative analysis of the aggregate diameter of hPDC aggregates treated with various concentrations of cytoskeletal inhibitors: Blebbistatin, Nocodazole, Y-27632, or Cytochalasin D (CytD). **f,** Semi-quantification of aggregate roundness for hPDC aggregates cultured for one or seven days with or without the cytoskeletal inhibitors blebbistatin, Y-27632 or CytD. **g,** Brightfield images of hPDC aggregates that were either untreated or treated with Y-27632 or blebbistatin for one week of culture. Scale bar: 100 µm. **h,** Fluorescence images of fixated monolayer hPDCs cells treated with iROCK (Y27632, 10 µM), Blebbistatin (10 µM) or CytD (5 µM) for one hour. The actin fibre organisation (phalloidin staining, white) and nuclei (DAPI, blue) are shown. Scale bar: 20 µm. **i,** Brightfield images of aggregates prior to CytD addition, after three days of culture with CytD, and following 15 days of recovery after the three days of treatment. Scale bar: 100 µm. j, Brightfield image showing unsuccessful aggregation of hPDCs seeded in microwells and let to assemble in the presence of CytD (5 µM) following six days of culture. k, Image-based semi-quantification of microtissue area. CytD (5 µM) was either directly added (direct), or added after successful aggregation (delayed). **l,** FluoCrowd analysis of hPDCs cells cultured in 2D vehicle control treated or CytD (5 µM) treated. Arrow indicated direction of volume change. Dots represent normalized median values. **m,** Maximum intensity projection fluorescence confocal images of F-actin in hPDCs cultured in either 2D or 3D prior to reseeding and imaging, Scale bar: 20 µm. **n,** Respective quantification of F-actin area and phalloidin staining intensity. **o,** Three-week chondrogenic differentiation of hPDC aggregates with or without low concentrations of CytD (0.5 µM), assessed by visualization of collagen matrix deposition (Pico Sirius Red staining) and **p**, respective image-based semi-quantification. n ≥ 16 microtissues per condition. Scale bars: 50 µm. The box plots in **a-f, k** give the mean ± 25-75%, with whiskers indicating the S.D.. Mean ± S.D. is shown in **n**. Statistical significance was assessed using Kruskal-Wallis ANOVA with Dunn post-hoc test (**n**).

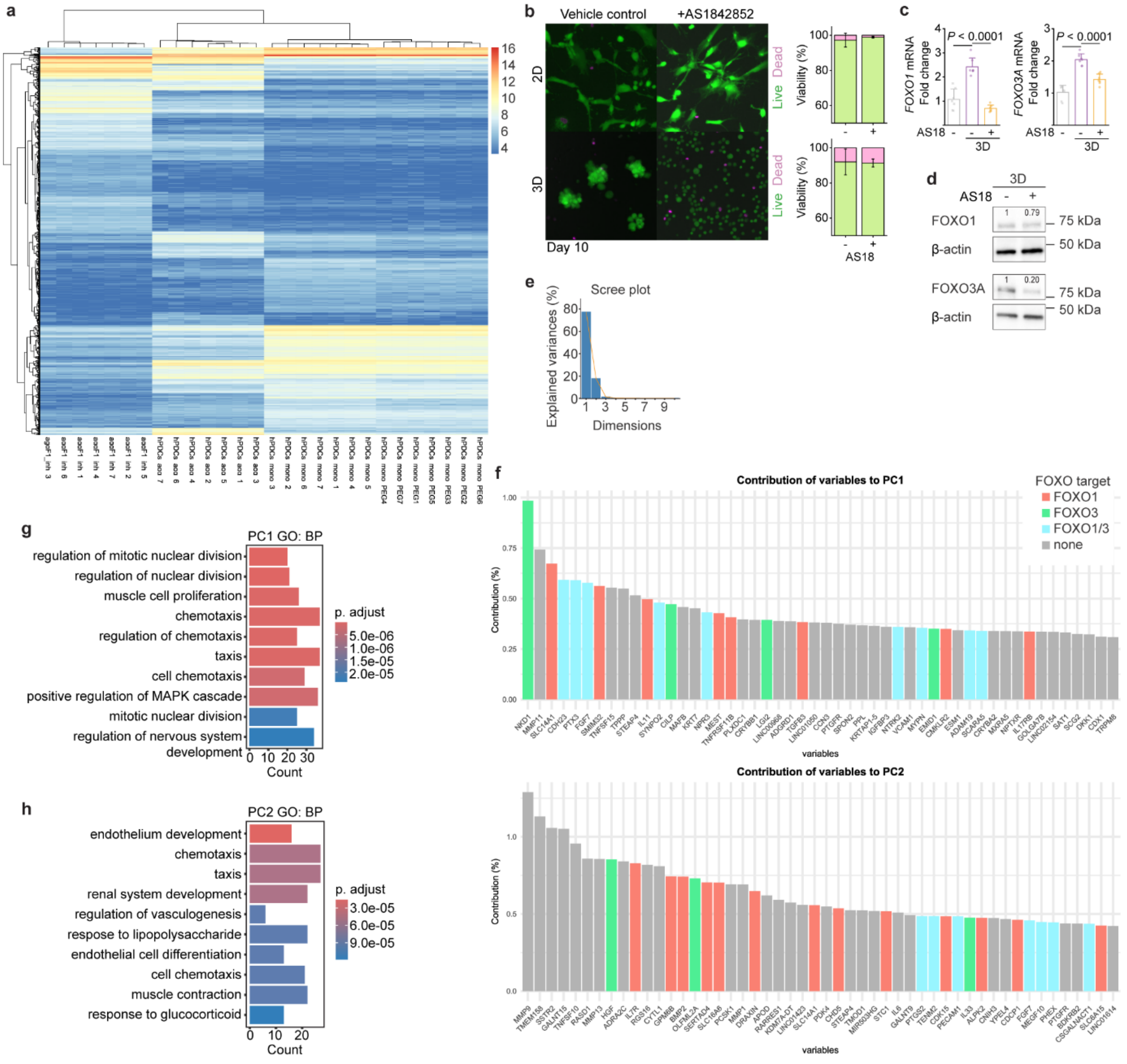


**Fig. S4 | a,** Heatmap of 1000 most variable genes by vst-expression shown for each sample. **b,** Fluorescence image of hPDCs showing cell viability cultured in the absence or presence of FOXO1 inhibitor (1 µM AS18) in either 2D or 3D for ten days, and respective quantification (n = 3 images per condition; n $\geq$ 290 cells (2D), n $\geq$ 1930 cells (3D)). Scale bar: 50 µm. **c,** FOXO1/3A mRNA levels in hPDCs cultured in 2D or 3D with or without FOXO1 inhibition (AS18; 1 µM). **d,** FOXO1 abundance in hPDCs cultured in 3D with or without FOXO1 inhibitor for three days. e, Scree plot, indicating that PC1 and PC2 explain most variance. **f,** Contribution of each gene on PC1 and PC2 expressed as eigenvalue and normalized to the sum of eigenvalues of all 1000 genes on PC1 and PC2. Shown are the top 20 contributors to PC1 and PC2. Colour coding indicates FOXO1/3 target genes. **g,** Pathway analysis results of PC1 contributors (> 0.05%) done with overrepresentation analysis on gene ontology (GO) terms for biological processes (BP). **h,** Pathway analysis results of PC2 contributors (> 0.05%) done with overrepresentation analysis on gene ontology (GO) terms for biological processes (BP). Mean ± S.D. is shown in **c**. Statistical significance was assessed using ANOVA with the Bonferroni post-hoc test (**c**). Protein abundance was normalized to β-actin abundance.

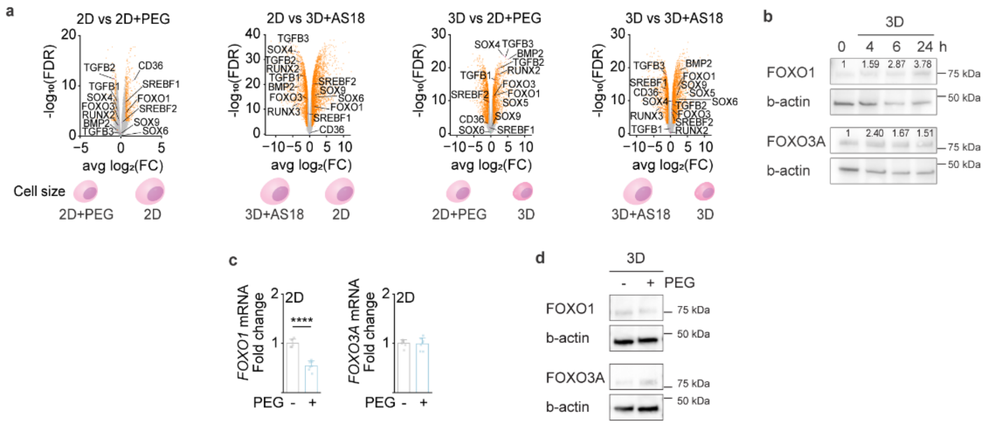


**Fig. S5 | a,** Volcano plots from RNAseq study comparing the four different experimental groups with each other. Cell volume changes between the compared conditions, are indicated in the little schematic below each plot. **b,** FOXO1/3 abundance over time within the first 24 hours following the seeding of hPDCs in agarose microwells. **c,** FOXO1/3 mRNA levels of hPDCs in response to 2% v/v PEG300 in 2D culture. **d,** FOXO1/3A abundance in hPDCs cultured in proliferation medium with or without 2% v/v PEG300 for three days. Statistics: Statistical significance was assessed using a two-sample t-test **(c).** Protein abundance was normalized to β-actin abundance.
